# Response of *Pseudomonas aeruginosa* to the innate immune system-derived oxidants hypochlorous acid and hypothiocyanous acid

**DOI:** 10.1101/2020.01.09.900639

**Authors:** Katie V. Farrant, Livia Spiga, Jane C. Davies, Huw D. Williams

## Abstract

*Pseudomonas aeruginosa* is a significant nosocomial pathogen and associated with lung infections in cystic fibrosis (CF). Once established, *P. aeruginosa* infections persist and are rarely eradicated despite the host immune cells producing antimicrobial oxidants, including hypochlorous acid (HOCl) and hypothiocyanous acid (HOSCN). There is limited knowledge as to how *P. aeruginosa* senses, responds to, and survives attack from HOCl and HOSCN, and the contribution of such responses to its success as a CF pathogen. We investigated the *P. aeruginosa* response to these oxidants by screening 707 transposon mutants, with mutations in regulatory genes, for altered growth following HOCl exposure. We identified regulators involved in antibiotic resistance, methionine biosynthesis and catabolite repression, and PA14_07340, the homologue of the *Escherichia coli* HOCl-sensor RclR (30% identical), that were required for HOCl survival. We have shown that RclR (PA14_07340) protects specifically against HOCl and HOSCN stress, and responds to both oxidants by upregulating expression of a putative peroxiredoxin, *rclX* (PA14_07355). While there was specificity in the transcriptional response to HOCl (231 genes upregulated) and HOSCN (105 genes upregulated) there was considerable overlap, with 74 genes upregulated by both oxidants. These included genes encoding the type III secretion system (T3SS), sulphur and taurine transport, and the MexEF-OprN efflux pump. RclR coordinated the transcriptional response to HOCl and HOSCN, including upregulation of pyocyanin biosynthesis genes, and in response to HOSCN alone RclR downregulated chaperone genes. These data indicate that the *P. aeruginosa* response to HOCl and HOSCN is multifaceted, with RclR playing an essential role.

---

*Pseudomonas aeruginosa* is a leading cause of nosocomial infections that are life threatening and difficult to treat (1). As an ESKAPE pathogen it is one of the bacteria that poses the greatest public health threat (2). Furthermore, *P. aeruginosa* is the major pathogen associated with chronic lung infections in patients suffering from cystic fibrosis (CF) (3,4). This genetic disease is caused by mutations in the cystic fibrosis transmembrane conductance regulator (CFTR), which results in abnormal ion and water transport across epithelial membranes, and leads to dehydrated airways and the establishment of respiratory infections (5). The immune response to infection is characterized by persistent inflammation that is ineffective at clearing infection and results in progressive lung tissue damage (3,4).

Host cells defend themselves against invading pathogens by the production of oxidants during the innate immune response, including the hypohalous acids: hypochlorous acid (HOCl), the active ingredient in bleach, and hypothiocyanous acid (HOSCN) (6). HOCl is a potent oxidant produced by neutrophils (7). The neutrophil oxidative burst that follows phagocytosis involves the reduction of O_2_ to superoxide (O_2_^−^) by the enzyme NADPH oxidase and the dismutation of O_2_^−^ to hydrogen peroxide (H_2_O_2_) (7). The haem enzyme myeloperoxidase (MPO) catalyses the formation of HOCl from the reaction of H_2_O_2_ with chloride (Cl^−^) (7). MPO can also mediate H_2_O_2_-oxidation of other halides including the pseudohalide thiocyanate (SCN^−^) to HOSCN, however its primary physiological substrate in neutrophils is proposed to be HOCl as the plasma concentration of Cl^−^ (100-140 mM) is much greater than SCN^−^ (20-100 μM) (6,8). At the epithelial cell surface of the lungs HOSCN is produced through the concerted action of the dual oxidase (DUOX) and lactoperoxidase (LPO) (9). The epithelial cell DUOX releases H_2_O_2_ into the airway surface liquid where LPO catalyses the reaction of H_2_O_2_ with SCN^−^ to form HOSCN (9). HOCl can target proteins, lipids, DNA and RNA; the most reactive targets of HOCl are thiol (S-H) groups of cysteine and methionine (7). Oxidation of cysteine by HOCl results in the formation of sulphenic acid, which can react with other cysteine thiols to form reversible disulphide bonds, and oxidation of methionine by HOCl results in formation of methionine sulphoxide (6). Similarly to HOCl, HOSCN reacts fastest with thiol groups of cysteine forming sulphenic acids or disulphide bonds (6).

Previous studies have identified transcriptional regulators that sense and respond to HOCl stress. These include the *Bacillus subtilis* transcriptional factors OhrR, HypR and PerR and the *E. coli* transcriptional factors NemR, HypT and RclR (10–14). All of the HOCl-sensing transcriptional regulators, apart from HypT, respond to HOCl through oxidation of cysteines, resulting in disulphide bond formation and activation or derepression of the regulators’ target genes (10–12,14–16). Instead, HypT responds to HOCl through oxidation of its methionine residues (17). HypT and RclR are the first described bacterial transcriptional regulators that are specific to sensing HOCl (13,14); the other regulators respond to a variety of stresses including reactive electrophile species and organic hydroperoxides (11,12,16,18–24). Identification of these HOCl-sensing transcriptional regulators revealed that bacteria have specific mechanisms for responding to HOCl stress, and do not rely solely on mechanisms used against other oxidants. The *E. coli* chaperones Hsp33 and RidA respond directly to HOCl via redox-switch mechanisms, which result in their activation and prevention of protein unfolding by HOCl-mediated oxidation of amino acid side chains (25,26). Virtually nothing is known about the response of bacteria to HOSCN, apart from a recent study from Groitl, et al. that investigated the transcriptional response of *P. aeruginosa* PA14 to HOSCN, as well as to HOCl and another hypohalous acid, hypobromous acid (HOBr) (27). A key finding of this study was that all three oxidants caused protein aggregation and *P. aeruginosa* responded by increasing levels of polyphosphate, which protected against protein aggregation and aided survival (27).

Here, we have used a combination of mutant screening and transcriptomics to identify and characterise mechanisms used by *P. aeruginosa* to survive HOCl and HOSCN challenge.

## Results

### Screening transposon mutants of P. aeruginosa regulatory genes for altered susceptibility to HOCl

We reasoned that targeting regulatory systems would be an effective way to uncover mechanisms used by *P. aeruginosa* to protect against HOCl stress. Therefore, we used gene ontology information from the *Pseudomonas* genome database (28) to compile a list of genes that were classed as having confirmed or predicted regulatory roles, and then determined whether there were mutants in these in the PA14 NR transposon mutant library (29). This resulted in a subset of 707 mutant strains (Table S1), which we screened for altered susceptibility when grown in the presence of 4.4 mM HOCl compared to WT. On screening, the most sensitive strains were identified as those that consistently failed to grow during the time of the assay or had an increased lag >3 hours compared to WT. Resistant strains were identified as those that consistently had a decreased lag >3 hours compared to WT. Fig. 1 shows an example of a 96-well screening plate with HOCl-sensitive and HOCl-resistant mutants labelled. Growth data for all plates screened is presented in the supporting information (Fig. S1). Identified HOCl-sensitive and HOCl-resistant mutants were rescreened and those showing visual altered HOCl-susceptibility are displayed in Fig. S2. Statistical analysis on these data revealed 16 mutants that were significantly HOCl-sensitive and 13 mutants that were significantly HOCl-resistant (Table 1 and 2).

**Figure 1.**
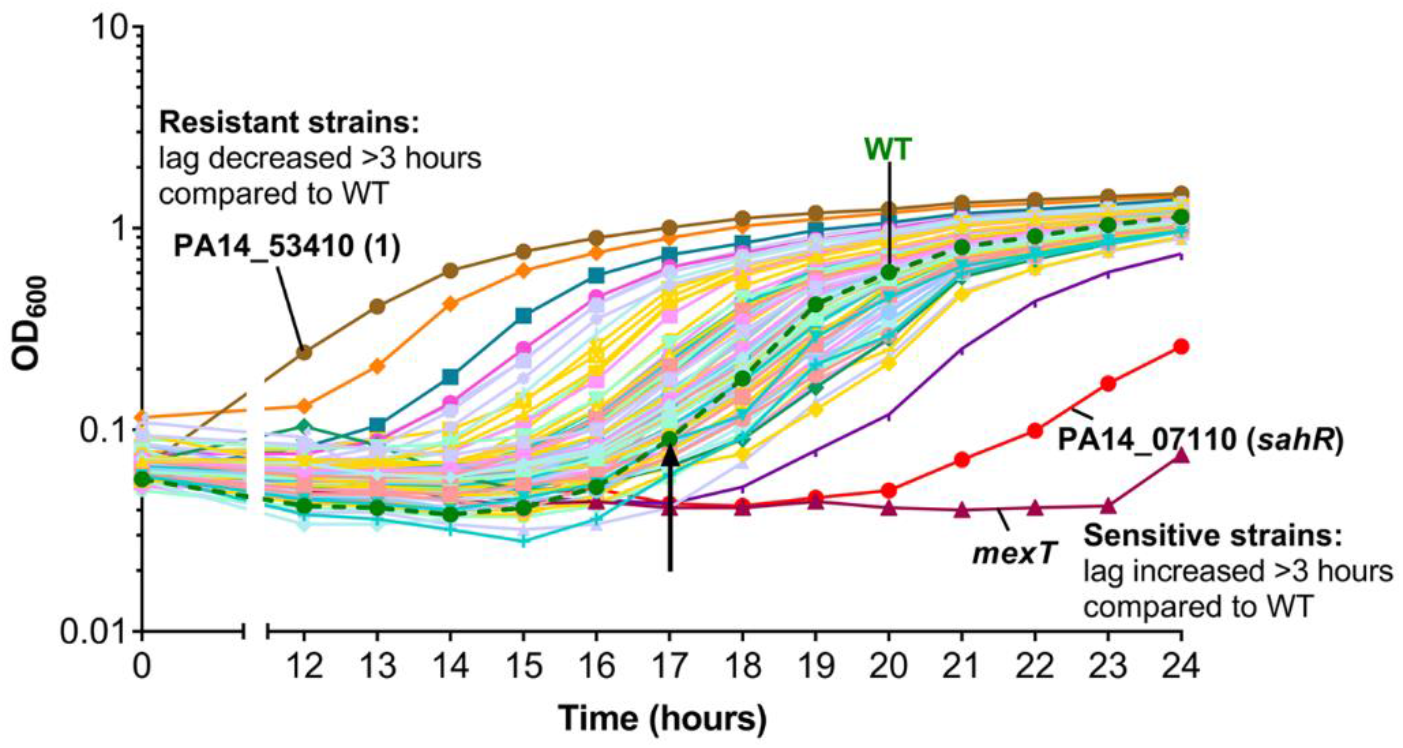
Identification of PA14 regulatory gene mutants with altered susceptibility to HOCl. Example screening plate with HOCl-sensitive (*mexT* and PA14_07110 (*sahR*)) and HOCl-resistant (PA14_53410 (1)) mutants labelled. Strains were grown in a 96-well format in LB medium with 4.4 mM HOCl and optical density (OD_600_) was recorded as a measure of growth. Mutant strains that had an increased lag >3 hours compared to WT were identified as HOCl-sensitive and strains with a decreased lag >3 hours compared to WT were identified as HOCl-resistant. Arrow represents the end of WT lag phase. Strains grown in the absence of HOCl showed no growth defects (data not shown).

**Table 1.**
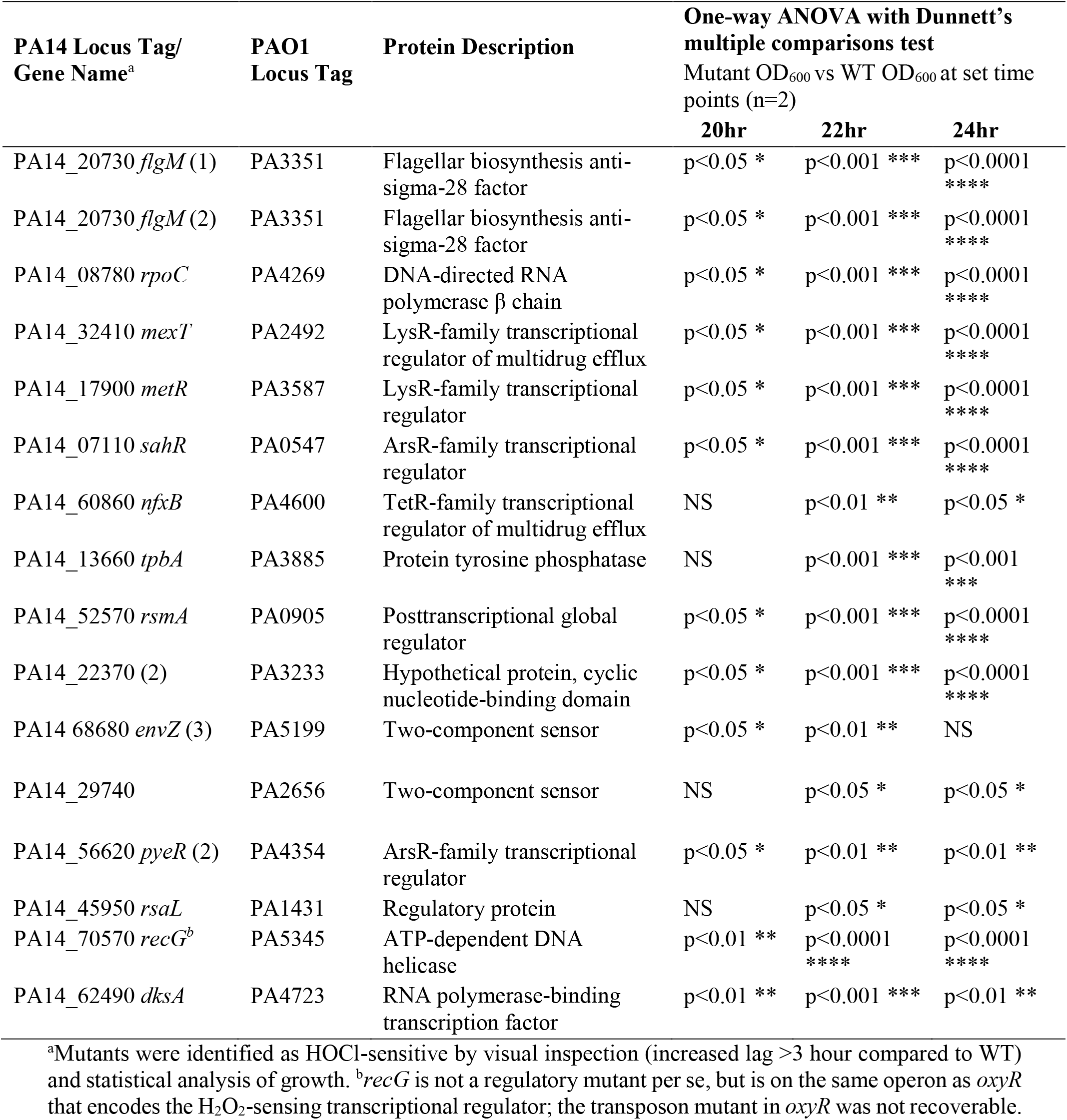
HOCl-sensitive regulatory gene mutants

**Table 2.** HOCl-resistant regulatory gene mutants

| PA14 Locus Tag/<br>Gene Name<sup>a</sup> | PAO1 Locus<br>Tag | Description | One-way ANOVA with Dunnett's multiple<br>comparisons test |  |  |  |
| --- | --- | --- | --- | --- | --- | --- |
|  |  |  | Mutant OD<sub>600</sub> vs WT OD<sub>600</sub> at set time points<br>(n=2) |  |  |  |
|  |  |  | 12hr | 14hr | 16hr | 18hr |
| PA14_10800 *ampR* | PA4109 | LysR-family<br>transcriptional regulator<br>of β-lactamase | p<0.0001<br>**** | p<0.0001<br>**** | p<0.0001<br>**** | p<0.001<br>*** |
| PA14_06950 (1) | PA0533 | LuxR-family<br>transcriptional regulator | p<0.0001<br>**** | p<0.0001<br>**** | p<0.0001<br>**** | p<0.001<br>*** |
| PA14_09760 | PA4185 | GntR-family<br>transcriptional regulator | NS | p<0.01 ** | p<0.05 * | NS |
| PA14_38040 *cmrA* | PA2047 | AraC-family<br>transcriptional regulator | p<0.0001<br>**** | p<0.0001<br>**** | p<0.0001<br>**** | p<0.0001<br>**** |
| PA14_44490 *anr* | PA1544 | Transcriptional regulator<br>of anaerobic metabolism | p<0.0001<br>**** | p<0.0001<br>**** | p<0.0001<br>**** | p<0.01 ** |
| PA14_27900 | PA2802 | GntR-family<br>transcriptional regulator | NS | p<0.01 ** | p<0.01 ** | p<0.05 * |
| PA14_68680 *envZ* (1) | PA5199 | Two-component sensor | p<0.0001<br>**** | p<0.0001<br>**** | p<0.0001<br>**** | p<0.01 ** |
| PA14_62530 *cbrA* (1) | PA4725 | Two-component sensor of<br>catabolic pathways | p<0.0001<br>**** | p<0.0001<br>**** | p<0.0001<br>**** | p<0.05 * |
| PA14_62530 *cbrA* (2) | PA4725 | Two-component sensor of<br>catabolic pathways | NS | p<0.05 * | p<0.05 * | NS |
| PA14_62540 *cbrB* | PA4726 | Two component response<br>regulator of catabolic<br>pathways | p<0.01 ** | p<0.001<br>*** | p<0.05 * | NS |
| PA14_50180 *fleR* (1) | PA1099 | Two-component response<br>regulator of flagellar<br>biosynthesis | NS | NS | p<0.05 * | NS |
| PA14_50200 *fleS* (2) | PA1098 | Two-component sensor of<br>flagellar biosynthesis | NS | p<0.0001<br>**** | p<0.01 ** | p<0.05 * |
| PA14_56620 *pyeR* (1) | PA4354 | ArsR-family<br>transcriptional regulator | p<0.0001<br>**** | p<0.0001<br>**** | p<0.0001<br>**** | p<0.001<br>*** |

<sup>a</sup>Mutants were identified as HOCl-resistant by visual inspection (decreased lag >3 hour compared to WT) and statistical analysis of growth.

### Links between antimicrobial resistance and HOCl susceptibility; the MexEF-OprN multidrug efflux pump protects P. aeruginosa from HOCl

A mutant in *mexT* was extremely sensitive to HOCl (Fig. 1 and Table 1). MexT is a LysR-type regulator that positively regulates expression of its adjacent operon *mexEF-oprN*, which encodes a multidrug efflux pump (30,31) (Fig. 2A). MexT regulates a number of genes (32,33), in addition to the *mexEF-oprN* operon, and so to determine whether loss of the efflux pump is associated with HOCl-sensitivity, transposon mutants of all three efflux pump genes were tested for altered HOCl susceptibility; they displayed increased sensitivity in order of *mexE* >*mexF*>*oprN* (Fig. 2A). This indicates that the MexEF-OprN efflux pump is needed for protection of *P. aeruginosa* against HOCl. Interestingly, an *ampR* mutant, which encodes a global regulator of over 500 genes and is a repressor of *mexEF-oprN* (34) was HOCl-resistant (Table 2, Fig. S1 and Fig. S2). MexT positively regulates 12 other genes (32) in addition to *mexEF-oprN* and *pyeR* (mentioned below); mutants in 8 of these were available for screening, and 4 displayed HOCl-sensitivity, albeit none were as sensitive as *mexE* (Table S2). Another multidrug efflux pump regulator, NfxB, was found to be required for protection against HOCl, as an *nfxB* mutant had increased HOCl-sensitivity (Table 1 and Fig. 2B). NfxB is a negative regulator of the *mexCD-oprJ* multidrug efflux operon (35) and therefore in the *nfxB* mutant overexpression of this efflux pump appears linked to HOCl-sensitivity. The lack of phenotypes of the *mexC* and *mexD* mutants is consistent with this, while the sensitivity of the *oprJ* mutant is unexpected (Fig. 2B). However, loss of function mutations in *nfxB* lead to global changes in *P. aeruginosa* physiology, which might impact HOCl-sensitivity (36).

**Figure 2.**
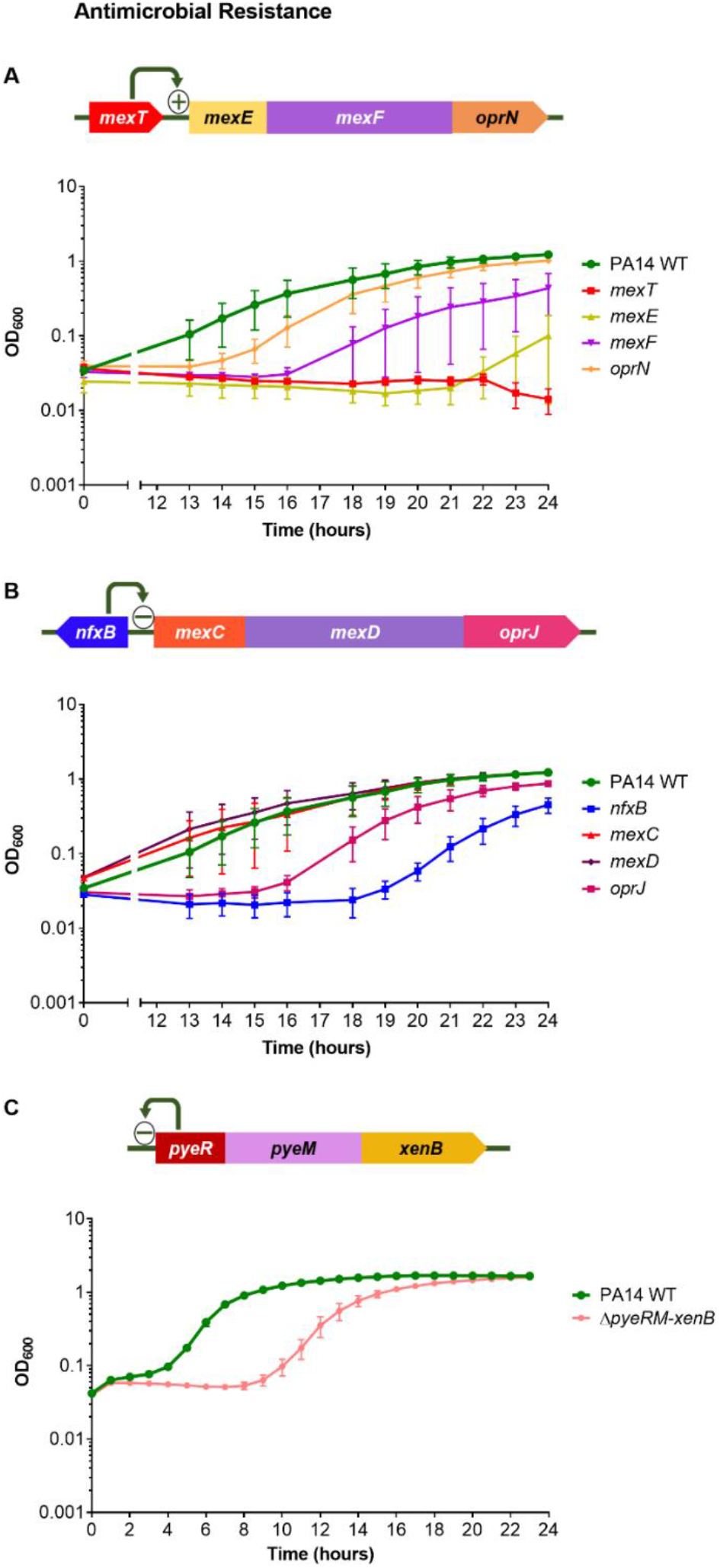
Mutants of genes involved in antimicrobial resistance displayed altered susceptibility to HOCl. (A) Gene arrangement of *mexT* and its adjacent *mexEF-oprN* operon. Growth of PA14 WT, *mexT*, *mexE*, *mexF*, and *oprN* strains in the presence of 4.4 mM HOCl. (B) Gene arrangement of *nfxB* and its adjacent *mexCD-oprJ* operon. Growth of PA14 WT, *nfxB*, *mexC*, *mexD* and *oprJ* strains in the presence of 4.4 mM HOCl. For *mexC* and *mexD*, 2 transposon mutants of each gene were available and showed similar phenotypes, therefore only 1 mutant of each is displayed. (C) Gene arrangement of the *pyeRM-xenB* operon; PyeR autorepresses the *pyeRM-xenB* operon (37). Growth of PA14 WT and Δ*pyeRM-xenB* strains in the presence of 5.1 mM HOCl. All graphs display three biological replicates, error bars represent standard error of the mean. Strains grown in the absence of HOCl showed no growth defects (data not shown). All transposon mutants and the in-frame deletion were confirmed by PCR.

MexT regulates the *pyeRM-xenB* operon, in which *pyeR* encodes an ArsR-family repressor, *pyeM* a major facilitator superfamily (MFS) transporter and *xenB* an uncharacterized xenobiotic reductase (37) (Fig. 2C). Transposon mutants in *pyeR* showed variable susceptibility to HOCl (Table 1 and 2, Fig. S2); therefore we constructed a whole operon in-frame deletion mutant, Δ*pyeRM-xenB*, which displayed HOCl-sensitivity (Fig. 2C). This suggests a role for this MexT-regulated operon in protecting *P. aeruginosa* against HOCl.

### Metabolic regulators are involved in protection against HOCl

A mutant in *metR*, which encodes a regulator homologous (40% identity) to *E. coli* MetR, a regulator of methionine biosynthesis genes (38), was HOCl-sensitive (Table 1 and Fig. 3A). A mutant in PA14_07110 (*sahR*) was also HOCl-sensitive (Fig. 1, Table 1 and Fig. 3A). This gene encodes an ortholog of SahR from *Desulfovibrio spp*, which is a transcriptional regulator of the S-adenosylmethionine (SAM) cycle genes that are involved in the recycling of methionine (39).

**Figure 3.**
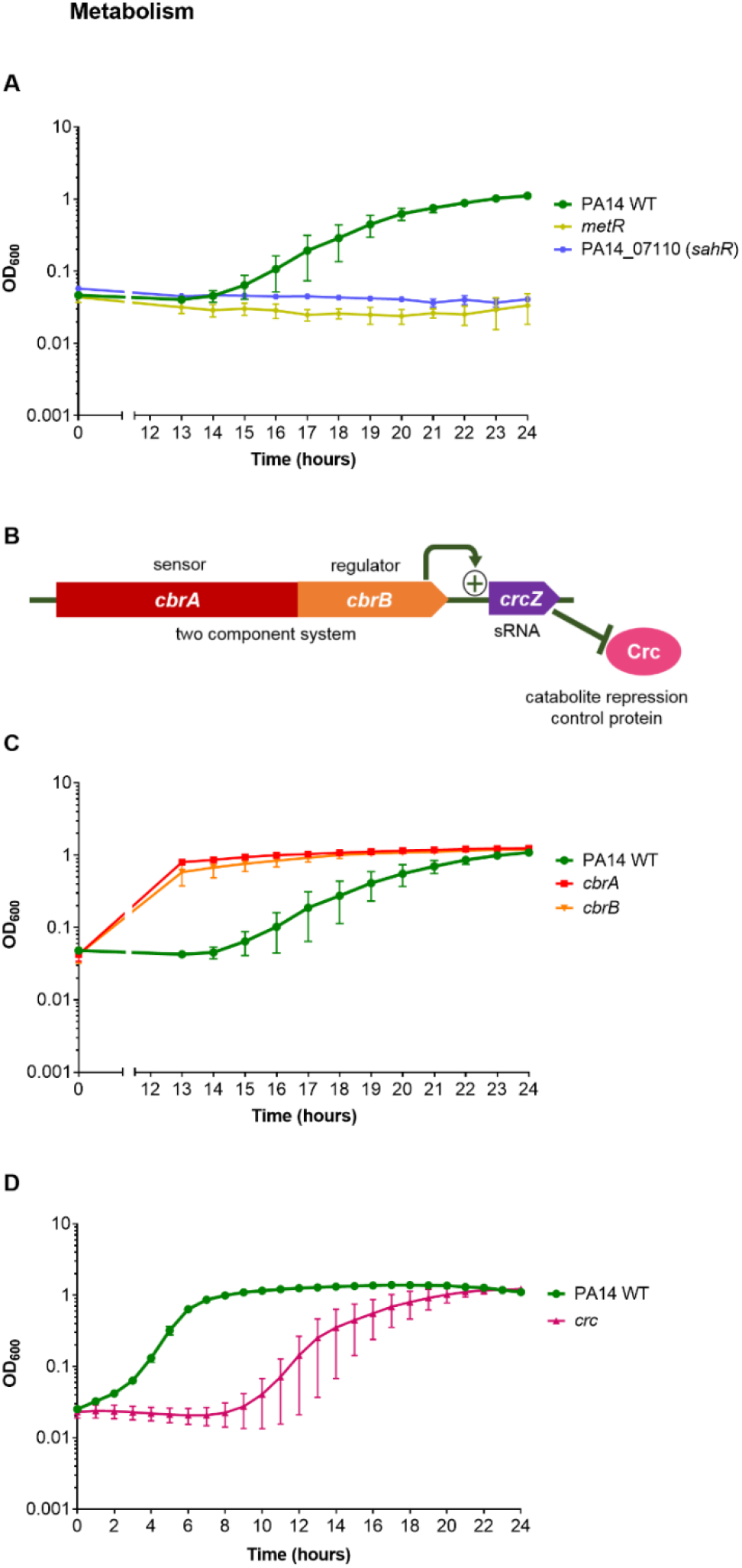
Mutants of genes involved in metabolism displayed altered susceptibility to HOCl. (A) Growth of PA14 WT, *metR* and PA14_07110 (*sahR*) strains in the presence of 4.4 mM HOCl. (B) Gene arrangement of the two-component system *cbrAB* that regulates expression of the sRNA CrcZ, which relieves Crc-mediated repression of target mRNAs. (C) Growth of PA14 WT, *cbrA* and *cbrB* strains and (D) PA14 WT, and *crc* strains, in the presence of 4.4 mM HOCl. For *cbrA*, 2 transposon mutants were available and showed similar phenotypes, therefore only 1 mutant is displayed. All graphs display three biological replicates, error bars represent standard error of the mean. Strains grown in the absence of HOCl showed no growth defects (data not shown). All transposon mutants were confirmed by PCR.

The CbrA/B two component system is involved in controlling the expression of carbon and nitrogen metabolism and carbon catabolite repression (40–42). CbrA/B acts through activation of the small RNA CrcZ, which sequesters the catabolite repression control protein Crc, relieving catabolite repression (42,43) (Fig. 3B). Mutants of *cbrA*/*B* were HOCl-resistant (Table 2 and Fig. 3C) and subsequently the *crc* transposon mutant was tested and found to be HOCl-sensitive (Fig. 3D). These phenotypes correlate with the opposing regulatory roles of CbrA/B and Crc.

### Regulators with putative roles in the oxidative stress response are involved in protection against HOCl

A mutant in the *algH* gene, which encodes a hypothetical protein that has been crystallised and predicted to be a potential redox-sensor, was HOCl-sensitive (Fig. 4A) (44). The *algH* gene is on a 4-gene operon that includes *gshB*, which encodes a glutathione synthetase; a *gshB* mutant was tested and displayed HOCl-sensitivity (Fig. 4A). A library mutant in *oxyR* encoding an H_2_O_2_-sensing transcriptional regulator (45) was non-viable, however a mutant was available for its adjacent gene a putative DNA repair enzyme, *recG* (45), and was HOCl-sensitive (Table 1 and Fig. 4B). The sensitivity of the *recG* mutant to HOCl was not due to disruption of *oxyR*, as the *oxyR* transcript was present in the *recG* mutant as confirmed by RT-PCR (data not shown). We constructed an in-frame deletion of *oxyR*, which also displayed HOCl-sensitivity (Fig. 4B). This suggests that the *oxyR-recG* operon is involved in protection of HOCl, as well as H_2_O_2_ (45).

**Figure 4.**
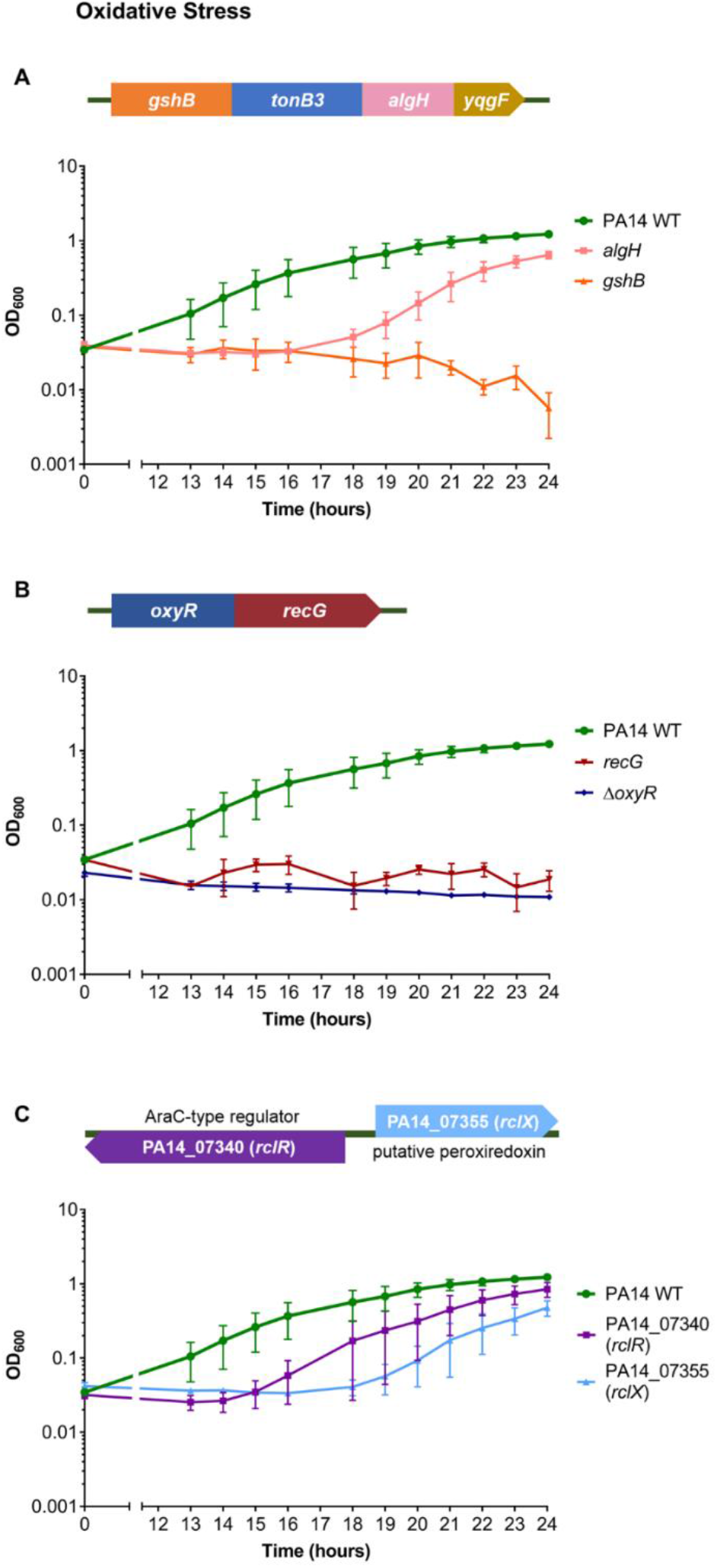
Mutants of genes involved in oxidative stress defence displayed altered susceptibility to HOCl. (A) Gene arrangement of the predicted 4-gene operon, *gshB*-*tonB3*-*algH*-*yqgF*. Growth of PA14 WT, *algH* and *gshB* strains in the presence of 4.4 mM HOCl. PA14 transposon mutants were not available for *yqgF* but were for *tonB3*, however they did not display altered HOCl susceptibility (data not shown). (B) Gene arrangement of the *oxyR-recG* operon. Growth of PA14 WT, *recG* and Δ*oxyR* strains in the presence of 4.4 mM HOCl. (C) Gene arrangement of PA14_07340 (*rclR*) and PA14_07355 (*rclX*). Growth of PA14 WT, PA14_07340 (*rclR*) and PA14_07355 (*rclX*) strains in the presence of 4.4 mM HOCl. All graphs display three biological replicates, error bars represent standard error of the mean (two biological replicates only for *recG*). Strains grown in the absence of HOCl showed no growth defects (data not shown). All transposon mutants and the in-frame deletion were confirmed by PCR.

*P. aeruginosa* mutants of homologues of other transcriptional regulators that have been found to respond to HOCl in bacteria, NemR (PA14_36300), HypT (PA14_71640) and OhrR (PA14_27230), did not display consistent HOCl-sensitive or resistant phenotypes in the screen (Fig. S3). However, a mutant in PA14_07340, the *P. aeruginosa* homologue of the *E. coli* reactive-chlorine specific transcriptional regulator, RclR (14) was identified as HOCl-sensitive (Fig. 4C). The *rclR* mutant was one of three regulatory mutants that showed consistent, reproducible sensitivity to HOCl (Fig. S2), but did not meet the stringent criteria for inclusion in Table 1. *P. aeruginosa rclR* (PA14_07340) is adjacent to a single-gene operon PA14_07355 that encodes a putative peroxiredoxin enzyme of the alkylhydroperoxidase (AhpD) family (Fig. 4C). A mutant in PA14_07355 was tested and displayed clear HOCl-sensitivity (Fig. 4C). Transposon mutants of *rclR* (PA0564) and the peroxiredoxin gene (PA0565) in the PAO1 strain background (46,47) were screened and appeared HOCl-sensitive (Fig. S4).

### P. aeruginosa RclR is specifically required for protection against HOCl and HOSCN

To examine whether *P. aeruginosa* RclR was required specifically for protection against HOCl, in-frame deletion mutants of *rclR* and the putative peroxiredoxin gene, which we named *rclX*, were constructed and tested for altered susceptibility against a range of reactive oxygen, nitrogen and electrophilic species. Initially the susceptibility of the mutants to HOCl was retested; Δ*rclR* and Δ*rclX* grew similarly to WT in the absence of oxidant (Fig. 5A), but neither mutant was able to grow in the presence of HOCl (Fig. 5B). Complementation of Δ*rclR* and Δ*rclX* with *rclR*-pUCP18 and *rclX*-pUCP18, respectively, restored growth in the presence of HOCl to that of WT (Fig. 5B). Exposure to other reactive oxygen species (H_2_O_2_, the superoxide generator methyl viologen, *tert*-butyl hydroperoxide (TBH)), reactive electrophilic species (*N*-ethylmaleimide (NEM), methylglyoxal, diamide) and the nitric oxide generator diethylamine NONOate (DEANO) did not alter the growth of Δ*rclR* compared to WT, indicating that *P. aeruginosa* RclR is not required for protection against these species (Fig. S5). Due to HOSCN being another physiologically relevant, thiol-reactive oxidant produced by the immune system (9), we were interested to determine whether Δ*rclR* and Δ*rclX* had altered sensitivity to this oxidant. Growth of Δ*rclR* and Δ*rclX* in the presence of 0.8 mM HOSCN revealed Δ*rclR* was sensitive to HOSCN, whereas Δ*rclX* was not at this concentration (Fig. 5C). Complementation of Δ*rclR* with *rclR*-pUCP18 restored growth in the presence of HOSCN (Fig. 5C). These data were supported by viability assays which showed HOCl inhibited growth of Δ*rclR* and Δ*rclX* (Fig. 5D), and HOSCN was bactericidal towards Δ*rclR* (Fig. 5E). When the HOSCN concentration was increased to 1 mM, Δ*rclX*, as well as Δ*rclR*, displayed sensitivity to this oxidant (Fig. S6). Therefore, both RclR and RclX are required for protection against HOCl and HOSCN.

**Figure 5.**
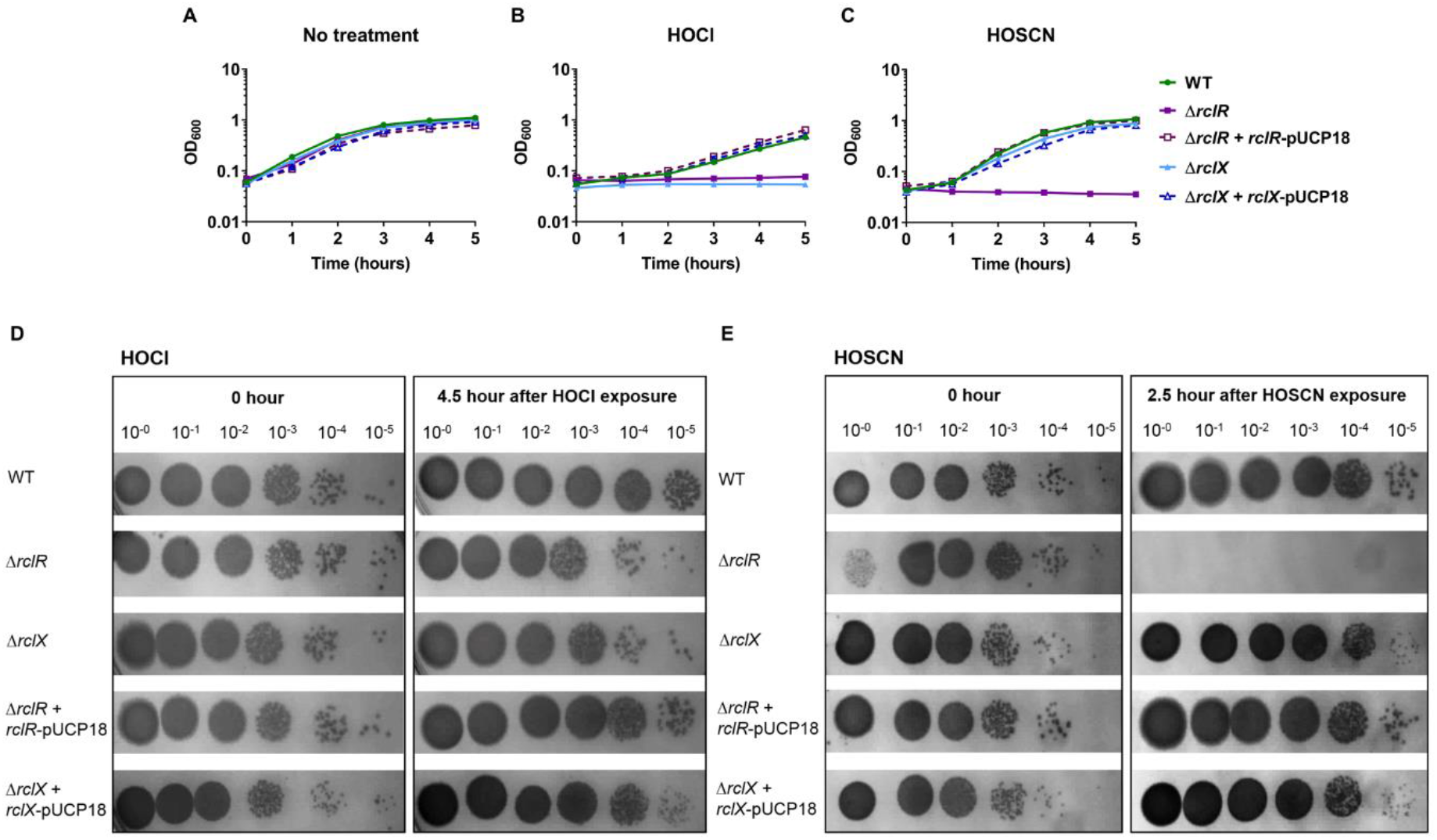
Mutants of *rclR* and its adjacent gene *rclX* are sensitive to HOCl, and *rclR* is sensitive to the epithelial-derived oxidant HOSCN. (A-C) Growth of PA14 WT, Δ*rclR*, Δ*rclR* + *rclR*-pUCP18, Δ*rclX*, Δ*rclX* + *rclX*-pUCP18 strains in (A) LB medium (B) LB medium with 4.4 mM HOCl or (C) LB medium with 0.8 mM HOSCN. Three biological replicates, error bars represent standard error of the mean. For HOSCN-sensitivity assays a control reaction of GO, glucose and KSCN was carried out, which resulted in production of H_2_O_2_ only; Δ*rclR* was not sensitive under these conditions confirming sensitivity is due to HOSCN production (data not shown). Addition of the empty pUCP18 vector to Δ*rclR* and Δ*rclX* had no effect on the susceptibility of these mutants to HOCl or HOSCN (data not shown). (D-E) Viability CFU assays, mid-exponential phase cultures of PA14 WT, Δ*rclR*, Δ*rclX,* Δ*rclR* + *rclR*-pUCP18 and Δ*rclX* + *rclX*-pUCP18 were incubated in LB medium containing (D) 4.4 mM HOCl or (E) 0.8 mM HOSCN. Samples were taken immediately after oxidant exposure and after 4.5 hours (HOCl) or 2.5 hours (HOSCN), and diluted in PBS and spot-titred onto LBA and incubated at 37°C overnight or until CFUs could be visualised. Images are representative of three biological replicates.

The sensitivity assays thus far were performed on planktonic cells in Luria-Bertani (LB) medium; to determine whether the findings remained consistent when grown under conditions relevant to the CF lung environment, sensitivity assays for Δ*rclR* and Δ*rclX* mutants were repeated in artificial sputum media (ASM), which mimics the sputum of CF patients (48,49). Both Δ*rclR* and Δ*rclX* displayed HOCl and HOSCN sensitivity when grown in ASM (Fig. S7), confirming the findings of the LB assays. Additionally, during chronic infections in the CF lung, *P. aeruginosa* persists in biofilms. To determine whether *P. aeruginosa* mutants display similar HOCl and HOSCN susceptibility when grown as biofilms, Δ*rclR* and Δ*rclX* mutants were grown in a suspended ASM biofilm assay in the absence or presence of HOCl and HOSCN (adapted from (49)). The WT, Δ*rclR* and Δ*rclX* strains formed biofilms in ASM in the absence of the oxidants (Fig. S8). However, in the presence of 3.5 mM HOCl and 0.53 mM HOSCN, both Δ*rclR* and Δ*rclX* were unable to grow and appeared sensitive to these oxidants, compared to WT that was able to form biofilms at these concentrations (Fig. S8). These data confirm the relevance of the impact of HOCl and HOSCN on *P. aeruginosa* in a more realistic CF environment.

### RclR regulates induction of rclX expression in response to HOCl and HOSCN stress

To determine whether RclR regulates expression of itself and/or *rclX* under HOCl and HOSCN stress, transcriptional *rclR-lacZ* and *rclX*-*lacZ* fusions were constructed and introduced into the WT and Δ*rclR* strains for *in vivo* β-galactosidase assays. The activity of *rclX-lacZ* increased following HOCl and HOSCN exposure in the WT strain, but not in the Δ*rclR* strain, suggesting expression of *rclX* increases following HOCl and HOSCN stress dependent on RclR regulation (Fig. 6A). No difference in the activity of *rclR-lacZ* was observed between WT and Δ*rclR* strains in the absence or presence of HOCl and HOSCN, suggesting RclR does not autoregulate its own expression and is constitutively expressed (Fig. 6A). We next examined if the activation of *rclX* in response to these oxidants occurred in clinical CF isolates of *P. aeruginosa*. The activity of *rclX-lacZ* in three CF isolates was measured in the absence or presence of HOCl and HOSCN. The *rclX* gene was significantly induced in response to both oxidants in all three clinical isolates compared to untreated controls (Fig. 6B). This confirmed that induction of *rclX* appears an important protective response to these oxidants across *P. aeruginosa* laboratory and clinical strains.

**Figure 6.**
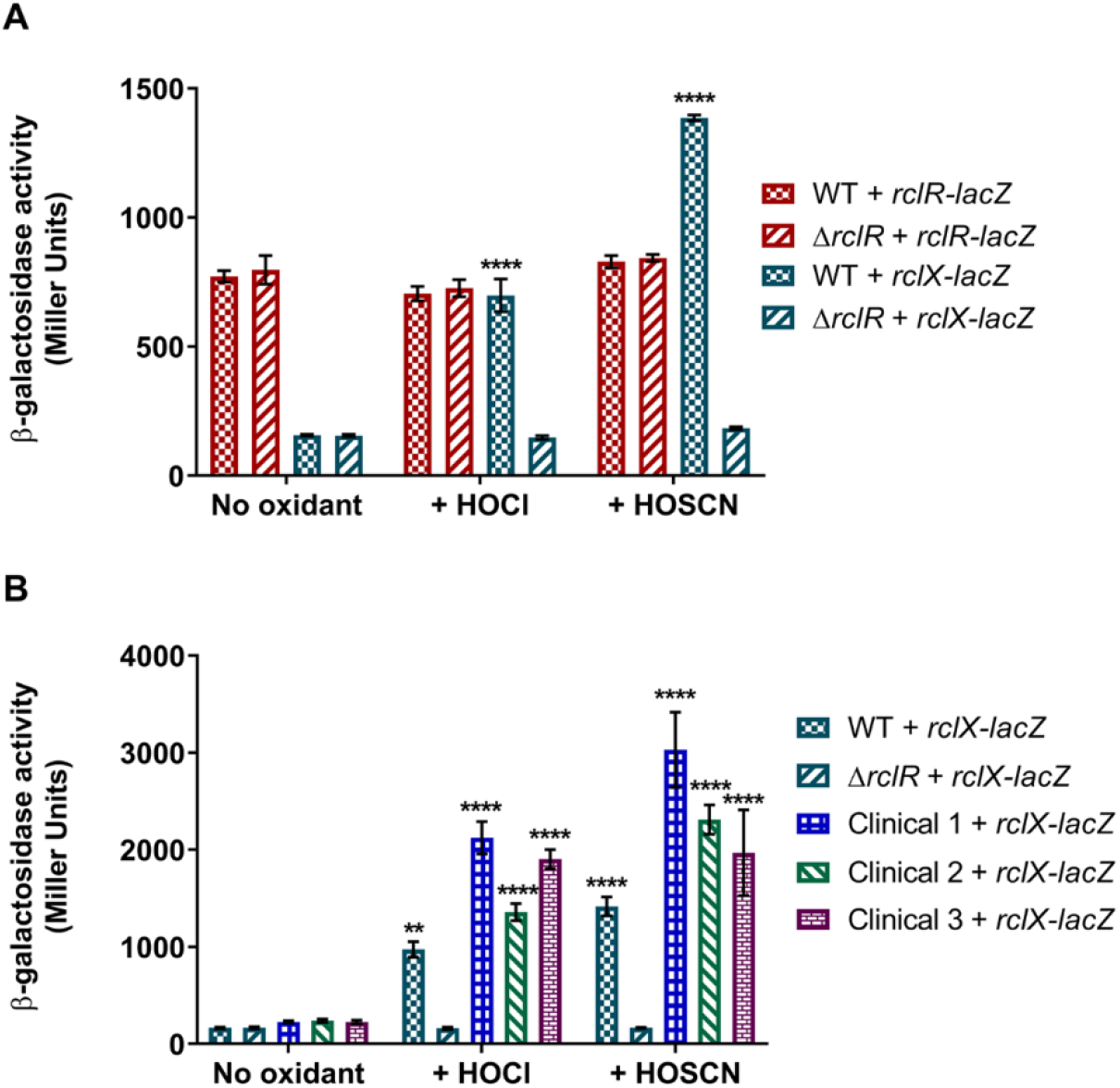
HOCl and HOSCN stress induces RclR-dependent regulation of *rclX* expression in PA14 *P. aeruginosa* and clinical CF isolates. (A) Activity of *rclR-lacZ* and *rclX-lacZ* transcriptional fusions in the PA14 WT and Δ*rclR* strains in the absence or presence of 2.2 mM HOCl or 0.8 mM HOSCN. (B) Activity of the *rclX-lacZ* transcriptional fusions in WT, Δ*rclR*, and clinical CF isolates 1, 2 and 3 in the absence or presence of 2.2 mM HOCl or 0.8 mM HOSCN. Strains were grown aerobically in LB medium until mid-exponential phase when HOCl or HOSCN was added and samples taken for β-galactosidase activity measurements at 30 minutes. PA14 WT and Δ*rclR* carrying the empty pMP220 vector expressed low β-galactosidase activity (~100 Miller Units) under all of the treatment conditions (data not shown). Error bars represent standard error of the mean of three biological replicates (six biological replicates for (B) untreated strains). Statistical analysis was performed using one-way ANOVA with Tukey’s multiple comparisons post-hoc test. ** indicates *p*<0.01 and **** indicates *p*<0.0001, compared to the same strain untreated (no oxidant).

To investigate the possible function of RclX, we performed homology modelling that fitted the RclX sequence with 32.1% sequence identity to the AhpD-like protein Lpg0406, from *Legionella pneumophila*, which belongs to the carboxymuconolactone decarboxylase family (50) (Fig. S9). The predicted structure was a homohexamer (Fig. S9A), formed from monomers with an AhpD-like fold containing six α-helices and a conserved catalytic CXXC motif (Fig. S9B). RclX was sequence aligned with Lpg0406, PA0269: another AhpD-like protein from *P. aeruginosa* (51), and the functionally characterised MtAhpD from *Mycobacterium tuberculous* (52–54) (Fig. S9C). Four out of the five catalytic residues of MtAhpD responsible for peroxidase activity (52,53) were conserved in RclX (Fig. S9C), suggesting a possible role for RclX in oxidant detoxification.

### Transcriptional changes in PA14 WT and ΔrclR in response to HOCl and HOSCN exposure

The response of *P. aeruginosa* to HOCl and HOSCN was further characterised by measuring changes in gene expression in the WT and Δ*rclR* strains. PA14 WT and Δ*rclR* exponential phase cultures were exposed to sub-lethal amounts of HOCl or HOSCN (2.2 or 0.8 mM, respectively) and gene expression analysed by RNA_seq_. Genes with a log2 fold change of >1.5 (2.9 fold change) were considered upregulated and genes with a log2 fold change of <−1.5 (0.35 fold change) were considered downregulated (Fig. 7). All transcriptomic data are presented in Data set S1 and are accessible through the GEO database under accession number GSE124385. HOCl upregulated 231 genes and downregulated 20 genes in the WT strain (Fig. 7A). HOSCN upregulated 105 genes and downregulated 16 genes in the WT strain (Fig. 7B). The importance of *rclX* in the *P. aeruginosa* response to HOCl and HOSCN was confirmed as it was strongly upregulated 72 and 448 fold change (>6 and >8 log2 fold change), respectively, and this was consistent with the gene fusion expression data (Fig. 7A+B and Fig 6). These data have been validated by qRT-PCR for a subset of 6 genes (including *rclX*), whose pattern of expression was confirmed (5 upregulated and 1 downregulated) (Fig. S10). For identification of genes whose expression is dependent on RclR, expression was compared in WT treated strains versus *rclR* treated strains. Therefore, in Fig.7C+D, genes displaying increased expression indicate genes positively-regulated by RclR and genes displaying decreased expression indicate genes negatively-regulated by RclR. RclR upregulated 42 genes and downregulated 7 genes in response to HOCl (Fig. 7C) and upregulated 132 genes and downregulated 213 genes in response to HOSCN (Fig. 7D). Regulation of *rclX* by RclR was confirmed as it was the gene most strongly upregulated by RclR in response to HOCl and HOSCN (Fig. 7C+D). Expression of *rclR* was not altered following exposure to HOCl or HOSCN, agreeing with our previous gene fusion data (Fig. 6A). Furthermore, the genes of homologues of other HOCl-responsive transcriptional regulators from *E. coli* and *B. subtilis*; NemR (PA14_36300), HypT (PA14_71640) and OhrR (PA14_27230) did not have altered expression in response to HOCl or HOSCN (Data set S1).

**Figure 7.**
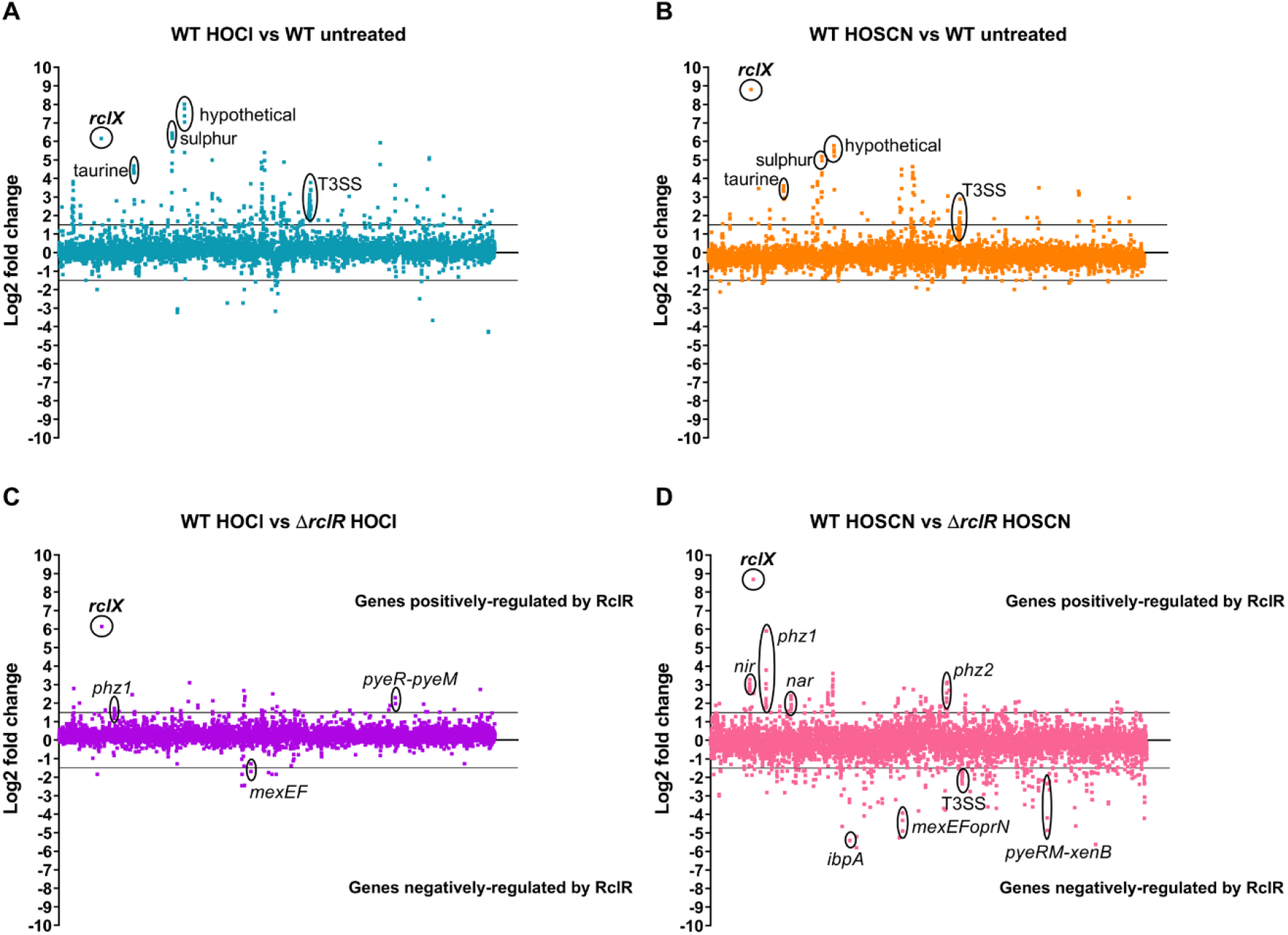
Relative changes in gene expression in PA14 WT and Δ*rclR* in response to HOCl or HOSCN exposure. Log2 fold change of gene expression plotted against the gene locus tag in order of genomic location for (A) WT+ HOCl vs. WT untreated, (B) WT+ HOSCN vs. WT untreated, (C) WT+ HOCl vs. Δ*rclR*+ HOCl or (D) WT+ HOSCN vs. Δ*rclR*+ HOSCN. Lines indicate cut off values at 1.5 or −1.5 log2 fold change. (A+B) Highlighted upregulated genes: *rclX*, hypothetical operon (PA14_21570-PA14_21580-PA14_21590-PA14_21600), sulphur transport genes (*ssuD*-PA14_19570-*ssuB*), taurine transport genes (PA14_12920-PA14_12940-PA14_12960), and T3SS genes. (C+D) In these plots positive log-fold change values indicate genes positively regulated by RclR and negative log-fold change values indicate genes negatively regulated by RclR. Highlighted genes positively regulated: *rclX*, pyocyanin biosynthesis operons *phz1* and *phz2*, denitrification *nir* and *nar* operons (HOSCN), and regulator *pyeR* and MFS transporter *pyeM* (HOCl); negatively regulated: *pyeRM-xenB* (HOSCN), drug efflux pump operon *mexEF-oprN*, T3SS genes (HOSCN), and heat shock protein *ibpA* (HOSCN).

The upregulated and downregulated genes were examined for overlap between the response to both oxidants and were categorised into functional groups based on assigned gene ontologies in the *Pseudomonas* genome database (28) (Fig. 8). Of the 105 genes upregulated in response to HOSCN, 70% (74 genes) were also upregulated in response to HOCl, indicating similarities between the bacterial responses to these two oxidants (Fig. 8A). The most highly upregulated group for both HOCl and HOSCN was non-coding RNA genes with 25% and 12.5% of the group being induced by these oxidants, respectively (Fig. 8B). Protein secretion/export apparatus genes and antibiotic resistance and susceptibility genes were the next highest induced groups (Fig. 8B). Of the 132 RclR-regulated genes whose expression was upregulated in response to HOSCN only 12 (9%) of those were also upregulated by HOCl (Fig. 8C), and of the 213 RclR-regulated genes downregulated following HOSCN exposure only 2 (1%) were downregulated after HOCl exposure (Fig. 8D). For RclR-regulated genes the most highly upregulated functional group following HOCl and HOSCN exposure was secreted factors, followed by energy metabolism (Fig. 8E). RclR downregulated a large number of genes in response to HOSCN, particularly those in the chaperone and heat shock proteins functional group (Fig. 8F).

**Figure 8.**
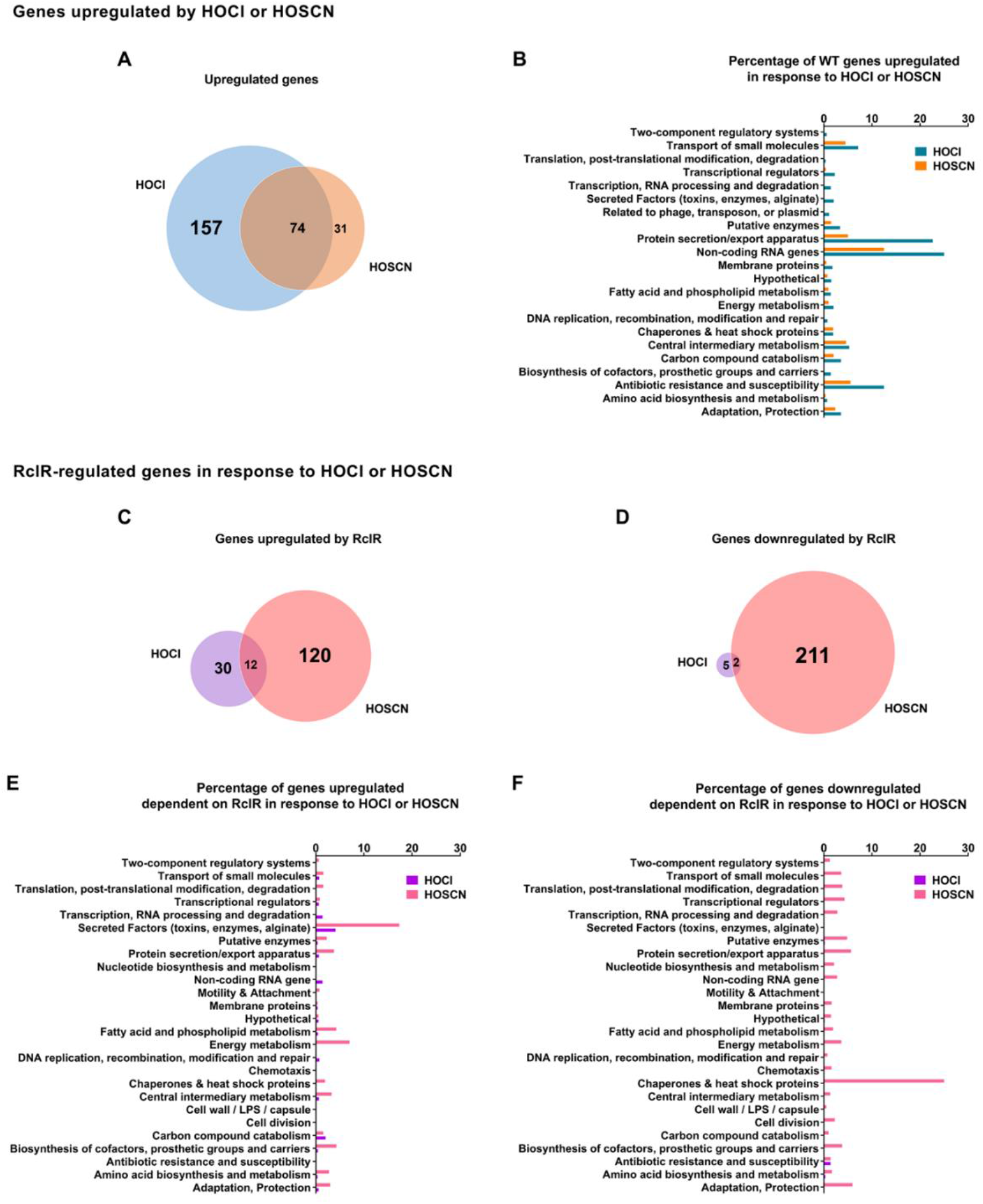
Numbers of upregulated or downregulated genes and functional categorisation of genes with altered expression in PA14 WT and Δ*rclR* after HOCl or HOSCN exposure. (A+B) Altered gene expression in the PA14 WT strain: (A) venn diagram of number of genes upregulated when comparing WT+ HOCl or HOSCN with WT untreated, (B) chart displaying percentage of upregulated genes in different functional groups. Only a small number of genes were downregulated in WT after HOCl (20) or HOSCN (16) exposure and are therefore not displayed. (C-F) Altered gene expression in the Δ*rclR* strain: venn diagrams of number of genes (C) upregulated or (D) downregulated when comparing WT+ HOCl or HOSCN with Δ*rclR*+ HOCl or HOSCN and charts displaying percentage of (E) upregulated or (F) downregulated genes in different functional groups. Venn diagrams display number of genes >1.5 log2 fold change upregulated or downregulated. Charts display genes categorised into functional groups based on assigned gene ontologies in the *Pseudomonas* genome library and expressed as a percentage of the entire functional category in the PA14 genome.

### tRNA, T3SS, sulphur and taurine metabolism, and MexT-regulated efflux pump genes show increased expression following exposure to HOCl and HOSCN

Expression of categorised genes was displayed in heat maps to demonstrate differences in gene expression in response to HOCl (first column) or HOSCN (second column) and in which upregulated genes are displayed in red and downregulated in green (Fig. 9 and Fig. 10). Genes that are positively or negatively regulated by RclR are indicated by red or green, respectively, in response to HOCl (third column) or HOSCN (fourth column) (Fig. 9 and Fig. 10).

**Figure 9.**
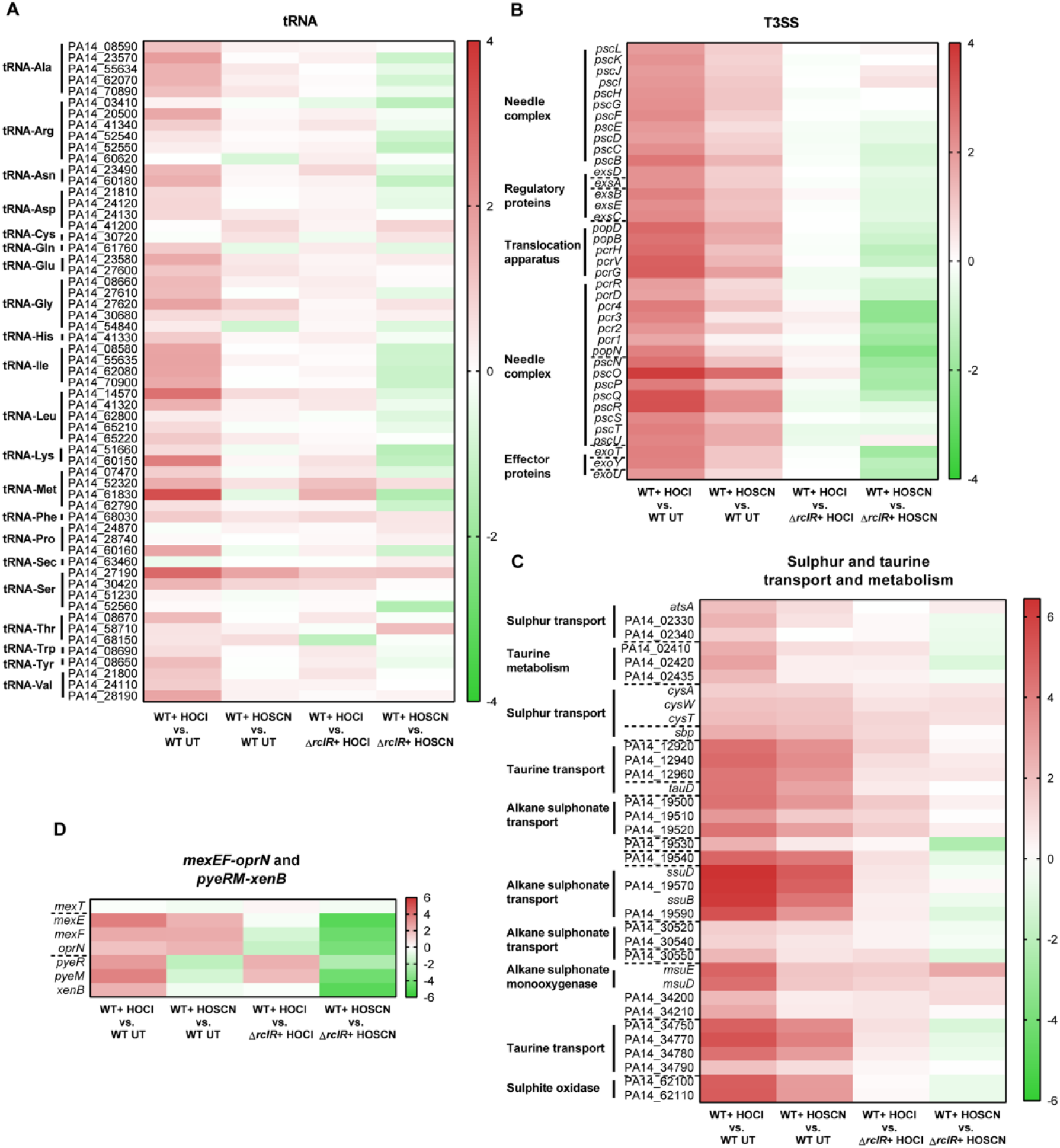
Heat maps displaying expression of tRNA, T3SS, sulphur and taurine transport and metabolism, and *mexEF-oprN* efflux pump and *pyeRM-xenB* genes in PA14 WT and Δ*rclR* after HOCl or HOSCN exposure. (A) tRNAs, (B) T3SS, (C) sulphur and taurine transport and metabolism and (D) *mexEF-oprN* and *pyeRM*-xenB. Expression of genes is colour coordinated from >6 log2 fold change (red) to −6 log2 fold change (green). The first column indicates HOCl responsive genes (log2 fold change of WT+ HOCl vs. WT untreated (UT)) and the second column indicates HOSCN responsive genes (log2 fold change of WT+ HOSCN vs. WT UT). The third column indicates RclR-regulated genes under HOCl stress (log2 fold change of WT+ HOCl vs. Δ*rclR*+ HOCl) and the fourth column indicates RclR-regulated genes under HOSCN stress (log2 fold change of WT+ HOSCN vs. Δ*rclR*+ HOSCN), and in these columns, genes that are positively or negatively regulated are indicated by red or green, respectively. Broken lines indicate the start and end of operons.

**Figure 10.**
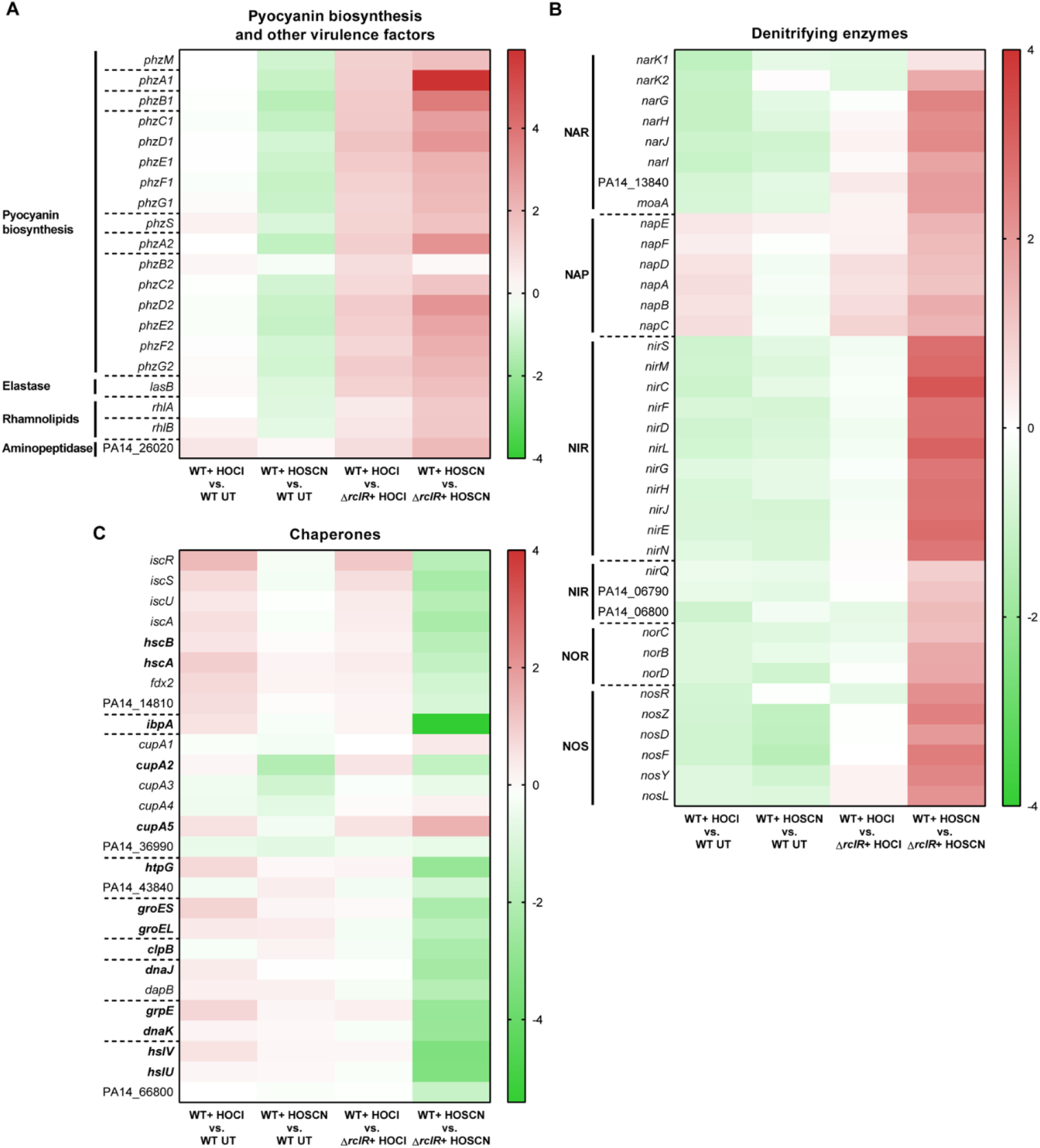
Heat maps displaying expression of pyocyanin biosynthesis, denitrifying enzyme, and chaperone and heat shock genes in PA14 WT and Δ*rclR* after HOCl or HOSCN exposure. (A) pyocyanin biosynthesis and other virulence factors, (B) denitrifying enzymes and (C) chaperones (genes in bold indicate the chaperones). Expression of genes is colour coordinated from >4 log2 fold change (red) to <−4 log2 fold change (green). The first column indicates HOCl responsive genes (log2 fold change of WT+ HOCl vs. WT UT) and the second column indicates HOSCN responsive genes (log2 fold change of WT+ HOSCN vs. WT UT). The third column indicates RclR-regulated genes under HOCl stress (log2 fold change of WT+ HOCl vs. Δ*rclR*+ HOCl) and the fourth column indicates RclR-regulated genes under HOSCN stress (log2 fold change of WT+ HOSCN vs. Δ*rclR*+ HOSCN), and in these columns, genes that are positively or negatively regulated are indicated by red or green, respectively. Broken lines indicate the start and end of operons.

The most highly upregulated genes in the WT strain in response to HOCl and HOSCN were from the non-coding RNAs functional group. HOCl exposure led to increased expression of tRNAs (Fig. 9A). In response to HOSCN, 5s and 16s rRNA were upregulated (Data set S1), but an rRNA depletion step was carried out prior to RNA_seq_ using the highly efficient Ribo-Zero kit (55) and less than 2% rRNA was recovered, therefore the physiological importance of this is unclear. All the genes in the T3SS operons were strongly induced in WT in response to both HOCl and HOSCN (Fig. 9B), while RclR appeared to downregulate a number of the T3SS genes in response to HOSCN (Fig. 9B). Around 40% of the genes in the transport of small molecules category that were upregulated in the WT response to HOCl and HOSCN were involved in sulphur or taurine metabolism (Fig. 9C). The most highly upregulated sulphur transport operon in response to HOCl and HOSCN was *ssuD*-PA14_19570*-ssuB*-PA14_19590, which encodes an alkanesulphonate transport system (Fig. 9C). Genes encoding other alkanesulphonate and sulphur transport systems, and the enzymes sulphite oxidase and alkanesulphonate monooxygenase, which are involved in sulphur metabolism, were also upregulated in response to HOCl and HOSCN (Fig. 9C). Taurine is the most abundant amino acid found in neutrophils (56) and genes encoding two taurine transport systems, PA14_12920-PA14_12940-PA14_12960-*tauD* and PA14_34750-PA14_34770-PA14_34780-PA14_34790, were upregulated in response to HOCl and HOSCN, and a three-gene operon (PA14_02410-PA14_02420-PA14_02435) encoding a putative taurine metabolism operon was also upregulated in response to HOCl (Fig. 9C).

The *mexEF-oprN* operon, which encodes the efflux pump previously found to be required for protection against HOCl (Fig. 2A), was strongly upregulated in response to HOCl and HOSCN exposure (Fig. 9D). The *mexT* regulator did not have altered gene expression (Fig. 9D). Surprisingly, these data indicated that RclR downregulates expression of the *mexEF-oprN* operon in response to HOCl and HOSCN (Fig. 9D). The MexT-regulated operon, *pyeRM-xenB*, which is also involved in protection against HOCl (Fig. 2C), was upregulated in response to HOCl (Fig. 9D), but was downregulated in response to HOSCN, and this appeared to be dependent on RclR regulation (Fig. 9D).

### RclR regulates expression of pyocyanin biosynthesis genes in response to HOCl and HOSCN, and denitrification and chaperone genes in response to HOSCN

The *phz* genes encode for biosynthesis of the virulence factor pyocyanin. While in the WT strain HOCl and HOSCN had no and little effect respectively on *phz* expression, the experiments with the Δ*rclR* mutant indicate that RclR is responsible for upregulating expression of the pyocyanin biosynthetic genes in response to both HOCl and HOSCN (Fig. 10A). In the absence of these oxidants, there was no difference between pyocyanin production in WT and Δ*rclR* strains (*P*-value of 0.8384, unpaired *t*-test) with mean values of 1.8 and 1.9 μg/ml, respectively (a negative control *phzM* transposon mutant produced no detectable pyocyanin). Similarly, the virulence factors elastase (*lasB*) and rhamnolipids (*rhlA* and *rhlB*) and an aminopeptidase (PA14_26020) did not display altered expression in response to HOCl or HOSCN in WT, but were induced by RclR in response to HOCl and HOSCN (Fig. 10A). In the WT strain HOCl and HOSCN had little effect on expression of genes required for anaerobic dissimilatory denitrification (Fig. 10B). However, in response to HOSCN, but not HOCl, RclR increased expression of 6 operons encoding the denitrifying enzymes: nitrate reductase (NAR), periplasmic nitrate reductase (NAP), nitrite reductase (NIR), nitric oxide reductase (NOR) and nitrous oxide reductase (NOS) (Fig. 10B). Expression of chaperone and heat shock genes in the WT strain were only slightly altered in response to HOCl and HOSCN (Fig. 10C). RclR did not regulate expression of these genes in response to HOCl, yet in response to HOSCN RclR is responsible for downregulating expression of 13 chaperone and heat shock genes *hcsB*, *hcsA*, *ibpA*, *cupA2*, *htpG*, *groES*, *groEL*, *clpB*, *dnaJ*, *grpE*, *dnaK, hslV*, and *hslU*, and upregulated expression of the chaperone gene *cupA5* (Fig. 10C).

## Discussion

During CF lung infections *P. aeruginosa* both directly damages the lung tissues and leads to the release of proinflammatory cytokines that result in massive neutrophil influx, which when activated release elastase, collagenase and antimicrobial oxidants that damage the lungs and impair bacterial clearance (3,4). One mechanism employed by the innate immune response to attack infecting bacteria is through the production and release of the antimicrobial oxidants HOCl and HOSCN by neutrophils and the airway epithelium, respectively (7,9).

In this study, we investigated how *P. aeruginosa* adapts to and protects itself against HOCl and HOSCN. This is of direct relevance to CF infections due to evidence of a compromised immune response in CF patients, including impaired HOCl and HOSCN formation (57–61), and evidence for HOSCN production being protective of lung function (62).

Our mutant screen identified regulators involved in antibiotic resistance, methionine biosynthesis, catabolite repression and the antioxidant response that were required for protecting *P. aeruginosa* against HOCl (Fig. 2–4). The RclR transcriptional regulator, the *P. aeruginosa* homologue of the *E. coli* RclR HOCl-specific sensor (14), was required specifically for protection of *P. aeruginosa* against HOCl and HOSCN and responded to the presence of these oxidants through activating expression of its adjacent gene, *rclX* (Fig. 5–7 and Fig. S5).

*E. coli* RclR (RclR_Ec_) also induces expression of its adjacent operon, *rclABC*, in the presence of HOCl (14). However, unlike *rclR*_*Ec*_ (14), *rclRPa* is constitutively expressed, is not upregulated by HOCl (nor HOSCN), and does not regulate its own expression (Fig. 6A). Our results are consistent with Groitl et al., who did not observe induction of *P. aeruginosa rclR* in response to HOCl, HOBr or HOSCN (27). The *rclX* gene regulated by RclR_Pa_ encodes a putative peroxiredoxin and is one of the most upregulated genes in response to HOCl and HOSCN exposure and is required for protection against both oxidants (Fig. 5–7 and Fig. S6). RclX is not homologous to any of the genes in the RclR_Ec_-regulated *rclABC* operon (14), indicating differing approaches to protection against HOCl between these gram-negative bacteria. Homology modelling and multiple sequence alignment support RclX being an AhpD-like protein, with 4 conserved catalytic residues including the CXXC motif (Fig. S9), which is required for peroxidase activity in the functionally characterised AhpD from *M. tuberculous* (52,53). Therefore, it is probable RclX acts as a detoxification enzyme in the protective response to HOCl and HOSCN. The sensitivity of Δ*rclR* and Δ*rclX* mutants to HOCl and HOSCN when grown in ASM in both planktonic and biofilm cultures (Fig. S7-8), together with HOCl and HOSCN dependent induction of *rclX* expression in clinical CF isolates of *P. aeruginosa* (Fig. 6B), indicate that the RclR-mediated response is important in CF clinical strains and may play a role during infection.

There was considerable overlap in the transcriptional response *of P. aeruginosa* to HOCl and HOSCN (Fig. 8A+B). The RclR regulon in response to HOSCN was larger than the regulon in response to HOCl (Fig. 8C+D). HOSCN caused RclR-dependent induction of pyocyanin biosynthesis genes and denitrification genes and repression of chaperone and heat shock genes (Fig. 10). Three other transcriptional regulators that were found to be HOCl-responsive in other bacteria: NemR (PA14_36300), HypT (PA14_71640) and OhrR (PA14_27230) (10,12,13), did not have altered expression in response to HOCl or HOSCN (Data set S1). In a previous transcriptome study, Groitl et al., reported that *P. aeruginosa* NemR was induced by HOCl and HOSCN, and *P. aeruginosa* HypT was induced by HOCl (27). These differences may be due to slight variations in methodological approaches, such as the use of different medium and oxidant concentrations. Nevertheless, we demonstrate consistencies in our results as *nemR*, *hypT*, and *ohrR* were not HOCl-responsive, nor were mutants in these genes HOCl-sensitive (Fig. S3).

Our study revealed that the bacterial response to HOCl and HOSCN is multifaceted and incorporates an array of mechanisms involved in metabolism, redox-sensing, macromolecule repair and detoxification, export of toxic compounds and virulence. Figure 11 summarises the physiological processes described here as playing roles in protecting *P. aeruginosa* against HOCl and HOSCN *in vitro* and, we hypothesise, in the CF lung during infection. Sulphur transporter and metabolism genes were upregulated following HOCl and HOSCN exposure, and methionine metabolism regulators SahR and MetR were required for survival against HOCl (Fig. 3A and Fig. 9C). During sulphur starvation bacteria are able to use alkanesulkphonates and taurine as a sulphur source (63–66). Therefore, it is plausible that *P. aeruginosa* requires increased uptake of alternative sulphur sources and careful control of methionine metabolism as a mechanism to maintain levels of sulphur-containing compounds that are under oxidative attack upon HOCl and HOSCN exposure. The upregulation of sulphur transport and metabolism, and methionine and cysteine biosynthesis genes in response to HOCl stress has been reported in other bacteria (67–70). The catabolite repression control system (42) was required for protection against HOCl indicating that appropriate regulation of carbon metabolism is important for survival against HOCl (Fig. 3B-D). RclR was a strong positive regulator of denitrification genes in response to HOSCN (Fig. 10B). As these experiments were carried out aerobically we would expect these Anr-regulated genes to be down regulated (71), therefore it indicates that there may be a physiological advantage to expressing these pathways, aerobically in the presence of HOSCN.

**Figure 11.**
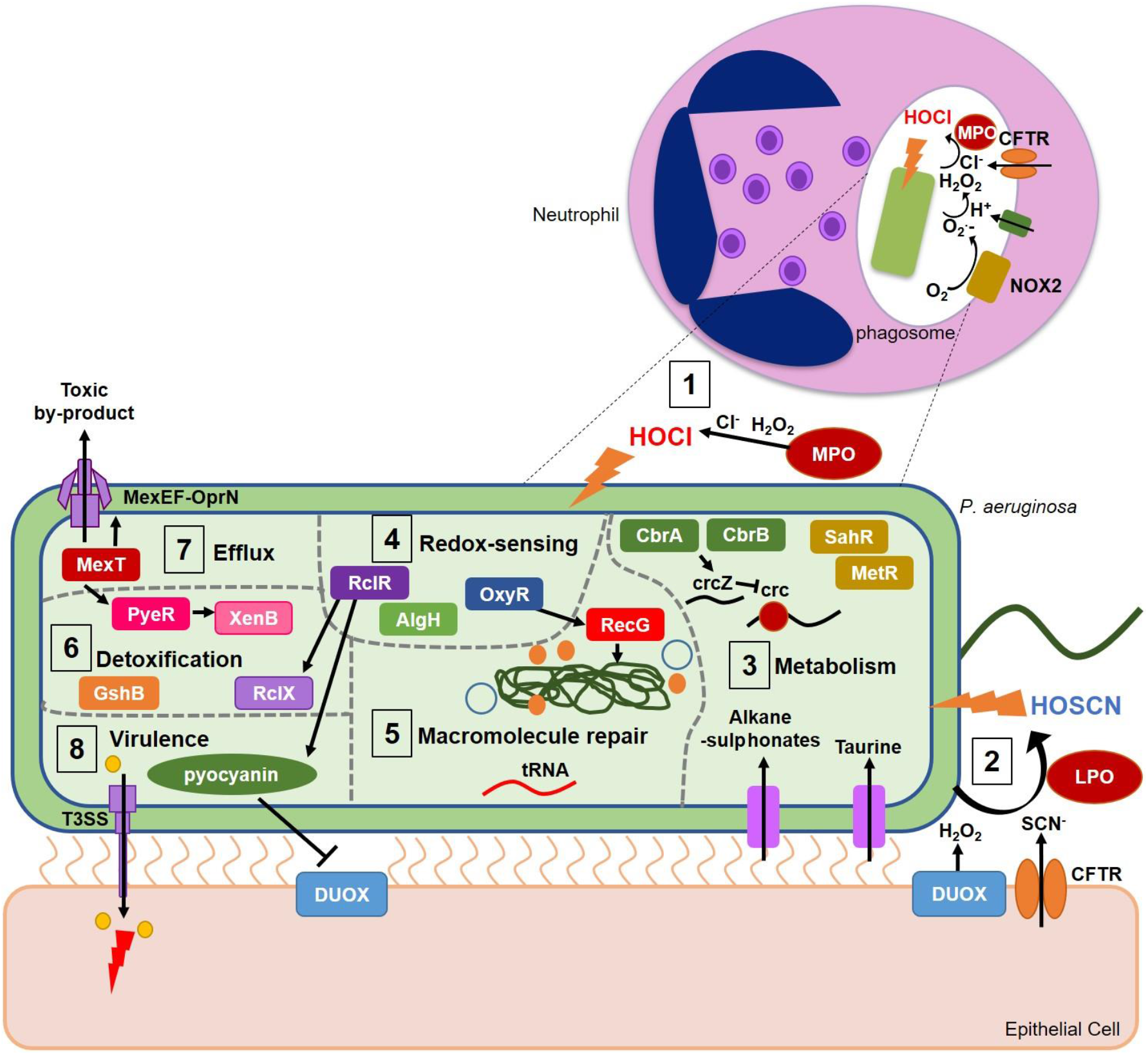
Model of the defence mechanisms used by *P. aeruginosa* in response to HOCl and HOSCN during infection. **1.** In the phagosome *P. aeruginosa* is exposed to HOCl produced by the MPO-catalysed reaction of Cl^−^ and H_2_O_2_ (7). HOCl is a potent oxidant that targets proteins, lipids DNA and RNA; its most reactive targets are sulphur-containing compounds and amines (6,7). **2.** The airway epithelial cells produce the oxidant HOSCN, via the LPO-catalysed reaction of H_2_O_2_ with SCN^−^ (9). HOSCN oxidises thiol groups of cysteine residues in bacterial proteins (6). **3.** *P. aeruginosa* responds to HOCl and HOSCN by upregulation of sulphur transport and metabolism genes; this may facilitate increased uptake of various alternative sulphur sources, including taurine, to replenish levels of sulphur depleted through the putative oxidative damage caused by HOCl and HOSCN. Methionine biosynthesis and metabolism regulators MetR and SahR are required to protect against HOCl, which we postulate is a response to oxidative disruption of the metabolism of the sulphur-containing amino acid methionine and the cellular need to maintain its levels. Catabolite repression by the Crc protein is required for protection against HOCl, indicating optimal regulation of metabolic flexibility aids survival against HOCl. **4.** The transcriptional regulator RclR, the H_2_O_2_-sensor OxyR (45) and the hypothetical protein AlgH (44), all have putative redox-sensing capabilities and were identified as being involved in protection against HOCl. **5.** Macromolecular repair mechanisms are implicated in the response to HOCl and HOSCN. Upregulation of tRNAs in response to HOCl may occur to replace oxidatively damaged tRNA. The DNA repair enzyme RecG is required for protection against HOCl. **6.** Putative bacterial detoxification enzymes are involved in the response to HOCl and HOSCN. The PyeR-regulated oxidoreductase XenB is induced in response to HOCl and is required for protection and the glutathione synthetase GshB is required for survival against HOCl. The peroxiredoxin RclX is positively regulated by RclR in response to HOCl and HOSCN and required for protection against both oxidants. **7.** Upregulation of the MexT-regulated *mexEF-oprN* operon, encoding a multidrug efflux pump (30,31) occurs in response to HOCl and HOSCN and is required for protection against HOCl. This highlights a link between oxidative stress and antibiotic resistance, and possibly indicates a role for the efflux pump in expelling toxic by-products of these oxidants. **8.** RclR is required for survival against HOCl and HOSCN and regulates a number of genes in response to both oxidants, including upregulation of pyocyanin biosynthesis genes. This may occur to maintain levels of pyocyanin that respond to the oxidative assault. Pyocyanin counterattacks the host immune response by DUOX inhibition and production of O_2_^−^, as well as inducing neutrophil apoptosis (86,87). Another virulence mechanism upregulated in response to HOCl and HOSCN is the T3SS, which is a needle-like machine that injects toxic effector proteins into host cells (88). Induction of the T3SS may occur as a mechanism to counterattack the host innate immune cells that are employing HOCl and HOSCN-mediated oxidative stress on the bacteria.

Genes involved in repairing macromolecules and detoxification of oxidants were important in the defence against HOCl and HOSCN. The H_2_O_2_-sensor OxyR and DNA repair enzyme RecG (45), were crucial for survival against HOCl (Fig. 4B). Also tRNA genes were upregulated in response to HOCl (Fig. 9A), perhaps to replace oxidatively damaged tRNA. RclR downregulated expression of 13 chaperone and heat shock genes, and upregulated expression of the chaperone gene *cupA5*, in response to HOSCN, but not HOCl (Fig. 10C). In the WT strain, expression of the chaperone gene *cupA2* was downregulated in the presence of HOSCN, but expression of the other genes was not altered in the presence of HOCl nor HOSCN (Fig. 10C). In contrast Groitl et al. found that HOCl and HOBr induced transcription of chaperone and heat shock genes, and together with HOSCN increased production of the chaperone polyphosphate, which protects *P. aeruginosa* against protein unfolding and aggregation caused by the oxidants (27).

We found detoxification enzymes, in addition to RclX, were required for survival against HOCl, including the *pyeRM-xenB* operon, which encoded the oxidoreductase XenB (37) (Fig. 2C). The glutathione synthetase GshB was required for protection against HOCl (Fig. 4A); this enzyme is involved in synthesising glutathione, which maintains the redox potential of the cell in response to oxidative stress (70). Previous transcriptome assays have highlighted the importance of repair and detoxifying enzymes in the response of bacteria to HOCl stress (67–69,72–75).

A striking finding of this work is a connection between the protective response to HOCl and HOSCN stress and antibiotic resistance. MexT and its regulated multidrug efflux pump MexEF-OprN was required for survival against HOCl (Fig. 2A) and the *mexEF-oprN* operon was strongly upregulated in response to both HOCl and HOSCN (Fig. 9D). While the fluoroquinolone, chloramphenicol and trimethoprim-exporting MexEF-OprN efflux pump is normally quiescent in wildtype cells it is highly induced in *nfxC* type phenotypic mutants (*mexT*, *mexS* or *mvaT*) in a MexT dependent manner (76). This leads to downregulation of expression of the quorum sensing-dependent efflux pump MexAB-OprM, and increased resistance to fluoroquinolones and chloramphenicol and hypersensitivity to β-lactams (77).

MexT has been shown previously to be a redox responsive regulator modulating the response to both electrophilic and nitrosative stress (78,79). Our data are consistent with recent findings (27) in extending the role of MexT and MexEF-OprN to HOCl and HOSCN stress protection and suggests that the cell has a pleiotropic need for induction of the MexEF-OprN efflux pump in response to a wide range of oxidative stress events. This leads to a model whereby MexEF-OprN is central in the response of *P. aeruginosa* to stresses affecting intracellular redox homeostasis including to HOCl and HOSCN production by the innate immune system.

Of direct relevance to CF lung infections is the repeated observation that mutations in *mexT* occur in persistently infecting *P. aeruginosa*, both loss of function, as well as, gain of function mutations (nfxC-type) (80–83). These would be predicted to cause *P. aeruginosa* to have reduced or enhanced ability, respectively, to defend itself against HOCl and HOSCN production by the innate immune response.

MexEF-OprN has previously been found to expel products other than antibiotics, and increased expression of *mexEF-oprN* is associated with decreased quorum sensing due to efflux of quorum sensing molecules, specifically the PQS precursors, HHQ and kynurenine (84,85). Therefore, one can speculate that MexEF-OprN might additionally expel toxic by-products of HOCl and HOSCN reactions. Consequently, MexEF-OprN induction has been linked to reduced levels of quorum sensing-dependent virulence traits including pyocyanin, HCN, elastase, rhamnolipids and T3SS (33).

Given that that HOCl/HOSCN induces MexEF-OprN induction, which would be expected to lead to PQS precursor efflux it might be expected that exposure to these oxidants also reduces the levels of PQS-dependent virulence factor production. While HOCl has no impact, HOSCN does downregulate pyocyanin biosynthesis, as well as the elastase encoding gene *lasB* and *rhlA* and *rhlB* that are involved in rhamnolipids synthesis (Fig. 10A). Surprisingly, RclR upregulates expression of pyocyanin biosynthesis genes in response to HOCl and especially HOSCN, as well as *lasB*, *rhlA* and *rhlB* (Fig. 10A), and so appears to be acting to maintain expression of these virulence factors genes under HOCl/HOSCN stress. In the case of pyocyanin this would facilitate a counterattack on the immune system’s oxidative assault, as pyocyanin is able to induce intracellular oxidative stress on the host airways by DUOX inhibition and production of O_2_^−^, and causes neutrophil apoptosis (86,87). Another virulence system, the T3SS (88), was induced by HOCl and HOSCN stress (Fig. 9B), which similarly may be postulated as an adaptive mechanism to counterattack the host innate immune cells that are employing HOCl and HOSCN-mediated oxidative stress on the bacteria.

In conclusion, our findings have identified a wide range of different genes involved in the bacterial response to HOCl and HOSCN stress, and provide the foundation from which further exploration of these mechanisms in protecting *P. aeruginosa* from these oxidants in the context of infection can be investigated.

## Experimental Procedures

### Bacterial strains, plasmids and growth conditions

All strains, plasmids and primers used in this study are listed in Table S1. Three anonymised clinical isolates of *P. aeruginosa* from airway secretions of infected CF patients were obtained from the Royal Brompton Hospital. PCR products and plasmids were sequenced by GENEWIZ, inc. Bacteria were routinely grown aerobically at 37^°^C, with shaking at 200 rpm or 700 rpm when in 96-well plates, in LB medium (5g/L NaCl, 5g/L yeast, 10 g/L tryptone) or on LBA plates (LB +1.5% w/v agar) supplemented with appropriate antibiotics. *P. aeruginosa* strains were isolated on *Pseudomonas* isolation agar (PIA) (Sigma) supplemented with 20% glycerol. ASM was prepared as described by Kirchner, et al. however all individual amino acids, apart from L-tryptophan, were replaced with casamino acids, as described in Sriramulu, et al. (48,49). The components of ASM per litre were: 4 g low molecular weight salmon sperm DNA, 5 g mucin from porcine stomach (type II), 4.75 g casamino acids, 0.25 g L-tryptophan, 5 g NaCl, 2.2 g KCl, 5.9 mg diethylenetriaminepentaacetic acid (DTPA), 5 ml egg yolk emulsion. The pH of the media was adjusted to 6.9 and filter sterilised (49) and stored at 4 °C. After filtration the media slowly turns cloudy, due to precipitation of the salts from the media, but this does not impact its quality. Pyocyanin was extracted from 20-hour cultures grown in LB and concentration expressed in μg/ml as described in (89).

### Construction of plasmids and mutant strains

Details of genes were obtained from The Pseudomonas Genome Database (28). In-frame deletions were constructed as previously described (90). Briefly, for construction of deletion mutants of the PA14 *rclR* and *rclX* genes, *pyeRM-xenB* operon, and *oxyR*, a ~500 bp upstream and ~500 bp downstream fragment of the genes was amplified by PCR using primer pairs 1F and 2R, and 3F and 4R (Table S1). Subsequent overlap PCR was performed using primers 1F and 4R (Table S1) to fuse the upstream and downstream fragments together to form a ‘mutator’ fragment that was purified and ligated into the pCR™-Blunt cloning vector (Invitrogen) for sequence-verification using M13F and M13R primers. The *rclR*, *rclX*, and *pyeRM-xenB* ‘mutator’ fragments were cloned into the BamHI site of the pKNG101 suicide vector and the *oxyR* ‘mutator’ fragment was cloned into the XbaI site of pKNG101, and all were transformed into the donor CC118λpir *E. coli* strain and plated onto LBA supplemented with 50 μg/ml streptomycin. The *rclR*-pKNG101, *rclX*-pKNG101, *pyeRM-xenB*-pKNG101, and *oxyR*-pKNG101 plasmids from the donor strain were introduced into the PA14 recipient strain by conjugation. The DH5α *E. coli* strain containing the helper plasmid pRK2013 was grown on LBA supplemented with 50 μg/ml kanamycin and the recipient PA14 strain was grown on LBA. Colonies of the donor, helper and recipient strains were mixed together on LBA and incubated overnight. PA14 with the *rclR*-pKNG101, *rclX*-pKNG101, *pyeRM-xenB*-pKNG101, and *oxyR*-pKNG101 plasmids integrated site-specifically into the chromosome as single-crossovers were isolated on PIA with 2000 μg/ml streptomycin. Subsequently, for sucrose counter-selection isolated colonies were streaked onto LBA without NaCl and with 10% (w/v) sucrose and incubated at room temperature for 48 hours to obtain double-crossover unmarked in-frame deletion mutants. For *oxyR* deletion 100 U of catalase from bovine liver per ml was added to the media. This is due to the *oxyR* mutants being unable to survive in LB, as components within the media are able to autoxidise to generate ~ 1.2 μM H_2_O_2_ min^−1^ (91). Colonies were isolated on LBA and deletion of *rclR*, *rclX*, *pyeRM-xenB*, and *oxyR* was confirmed by PCR and sequencing using 5F and 6R primers (Table S1). Subsequent initial overnight growth of Δ*oxyR* on LBA or in LB broth included 100 U/ml catalase from bovine liver. Complement plasmids, *rclR*-pUCP18 and *rclX*-pUCP18, were constructed by PCR amplification of the promoter region and ORF of PA14 *rclR* and *rclX* using primers in Table S1 and subsequent cloning into the HindIII/XbaI sites of the pUCP18 shuttle vector. The plasmids were transformed into DH5α *E. coli* and grown on LBA with 100 μg/ml ampicillin, prior to sequence verification using M13F and M13R primers (Table S1) and transformation into PA14 Δ*rclR* or Δ*rclX* mutant strains. The *rclR-lacZ* and *rclX-lacZ* plasmids were constructed by PCR amplification of the upstream promoter DNA (117 bp) of *rclR* and *rclX* using primers in Table S1 and ligation into pCR™-Blunt (Invitrogen) for sequence-verification using M13F and M13R primers. The promoter DNA of *rclX* and *rclR* was cloned into the EcoRI/PstI sites of the pMP220 promoter-probe vector (containing the *lacZ* gene) and transformed into DH5α *E. coli* and grown on LBA with 25 μg/ml tetracycline. Plasmids were sequence verified using primers Mid-PrclR or Mid-PrclX with Mid-lacZ (Table S1) prior to conjugation into PA14 WT and Δ*rclR*. Additionally, the *rclX-lacZ* plasmid was conjugated into three clinical CF isolates.

### Preparation of HOCl and HOSCN solutions

Sodium hypochlorite solution with 10-15 % available chlorine (Sigma) was diluted directly into LB or ASM to prepare the final HOCl concentrations required for assays. The concentration of HOCl was determined by diluting sodium hypochlorite into 40 mM potassium phosphate buffer (pH 7.5) with 10 mM sodium hydroxide and measuring the absorbance at 292 nM (ɛ 350 M^−1^ cm^−1^) (27,92). HOSCN was generated enzymatically by combining glucose, glucose oxidase (GO), potassium thiocyanate (KSCN) and LPO. GO catalyses the reaction of glucose with O_2_ to form H_2_O_2_, which reacts with KSCN, catalysed by LPO, to form HOSCN. Stock concentrations of the enzymes 200 U/ml GO (Sigma) and 1000 U/ml LPO (Sigma) were prepared in 50% PBS and 50% glycerol solution and stored at −20^°^C. Stock concentrations of 4% w/v glucose and 750 mM KSCN (Fluka, 8M) were prepared in H_2_O and stored at 4^°^C. Varying the amounts of glucose altered the amount of HOSCN produced and a final sublethal concentration of 0.01% glucose was selected for all assays performed in LB (apart from Fig. S6, for which a final amount of 0.02% glucose was used). The final concentrations of LPO, GO and KSCN were based on those used in (93). For the assays all components were diluted directly into LB or ASM to final concentrations of 0.01% glucose (0.02% glucose for Fig. S6), 0.5 U/ml GO, 0.75 mM KSCN and 3 U/ml LPO. A control assay without LPO that would only result in H_2_O_2_ generation (0.01% glucose, 0.5 U/ml GO and 0.75 mM KSCN) was performed to examine the effect of H_2_O_2_ (data not shown). The concentration of HOSCN was determined by reaction with 5-thio-2-nitrobenzoic acid (TNB), as described previously (27,94). Briefly, 10 mM 5,5’-dithiobis (2-nitrobenzoic acid) (Thermo Scientific) prepared in 40 mM potassium phosphate buffer (pH 7.5) was hydrolysed by addition of sodium hydroxide to generate TNB. The TNB concentration was determined by measuring absorbance at 412 nM (ε 14150 M^−1^ cm^−1^). HOSCN (0.01% glucose (0.02% glucose for Fig. S6), 0.5 U/ml GO, 0.75 mM KSCN and 3 U/ml LPO) prepared in 40 mM potassium phosphate buffer (pH 7.5) was diluted 1:1 in TNB and the loss of absorption at 412 nM was measured after 20 minutes. The concentration of HOSCN was determined from the concentration of TNB consumed during the reaction (stoichiometry 2:1, TNB: HOSCN). A concentration of 0.8 mM HOSCN was produced from 0.01% glucose and 1 mM HOSCN from 0.02% glucose. For the ASM assays 0.8 mM HOSCN was produced, as described above, which was then diluted in ASM to give the final HOSCN concentrations used.

### HOCl and HOSCN susceptibility and viability assays

A total of 707 strains with mutations in genes with regulatory functions were selected from the PA14 non-redundant single transposon mutant library (29) (Table S1). The strains were grown overnight in LB medium with 15 μg/ml gentamicin, alongside a PA14 WT control grown in LB medium, in 2.2ml 96-well deep well plates (VWR). For the HOCl screen, overnight cultures were subcultured 1:20 into LB medium with 15 μg/ml gentamicin in 96-well microtitre plates (Falcon) and grown for 3 hours. Cultures were subcultured again 1:20 (optical density (OD) 0.05 ± 0.03) in LB medium or LB medium containing 4.4 mM HOCl in 96-well microtitre plates (Falcon) and grown for up to 24 hours with OD_600_ recorded as a measure of growth at hourly time points. The concentration of HOCl used was selected from a preliminary experiment, which identified 4.4 mM HOCl as being sub-lethal to PA14 WT; causing a lag in growth but not complete inhibition (data not shown). However, for the assays that tested the HOCl susceptibility of Δ*pyeRM-xenB*, 5.1 mM HOCl was used. For PA14 WT, Δ*rclR* and Δ*rclX* mutant strains, and Δ*rclR*+ *rclR*-pUCP18 and Δ*rclX*+ *rclX*-pUCP18 complemented strains, overnight cultures were prepared by growing the strains in LB medium or LB medium with 500 μg/ml carbenicillin for the pUCP18 complemented strains. Overnight cultures were subcultured 1:20 in LB medium and grown for 3 hours, prior to subculturing again 1:20 (OD 0.05 ± 0.02) into LB medium or LB medium with 4.4 mM HOCl or 0.8 mM HOSCN and following growth by recording OD_600_. For the HOCl and HOSCN susceptibility assays in ASM media, the PA14 WT, Δ*rclR* and Δ*rclX* mutant strains were grown as above. The only difference was the ASM was filtrated again prior to adding HOCl at a final concentration of 2.5 or 3.1 mM, and adding components to make 0.8 mM HOSCN, which was diluted to final concentrations of 0.35 mM or 0.53 mM. Lower concentrations were required in the ASM assays as the bacteria had increased susceptible to oxidants in this media. For HOCl and HOSCN viability assays, cultures were sampled at 0 hour and 2.5 hour or 4.5 hour after HOSCN or HOCl exposure, respectively, and 10-fold serial dilutions in PBS were prepared and dilutions were drop plated on LBA. For other chemical susceptibility assays, PA14 WT, Δ*rclR* and Δ*rclX* strains were grown as before, but in LB medium containing 0.125 mM NEM, 5 mM diamide, 2.5 mM methylglyoxal, 5 mM H_2_O_2_, 0.5 mM TBH, 1 mM methyl viologen or 10 mM DEANO. The concentrations of chemicals used were determined from preliminary experiments that tested a range of concentrations to identify those that caused a lag, but not inhibition, in growth of PA14 WT (data not shown). All chemicals were from Sigma apart from TBH that was from EMD Millipore. For ASM biofilm assays we adapted the method from (49). Overnight cultures of PA14 WT, Δ*rclR* and Δ*rclX*, were subcultured 1:50 in LB and grown for 3 hours, prior to subculturing again 1:50 into 2ml ASM without or with HOCl (3.1 or 3.5 mM) and HOSCN (0.43 mM or 0.53 mM) in 24-well tissue culture treated plate (Falcon), which was incubated at 37 °C, low shaking, for 3 days to allow biofilms to form, and then left at room temperature, static, up to 7 days.

### RNA extraction and RNA_seq_ analysis

Exponential phase PA14 WT and Δ*rclR* cultures were subcultured 1:50 in LB (OD 0.05 ± 0.02) and grown for 2.5 hours prior to adding HOCl to a final concentration of 2.2 mM or HOSCN to a final concentration of 0.8 mM, or no treatment. The concentrations of oxidants used were chosen due to causing a 30-120 minute lag in growth of exponential phase cultures. WT and Δ*rclR* cultures were incubated for a further 20 minutes and then cells were collected and centrifuged at 13000 *x g* for 5 minutes. The pellets were washed twice with PBS and RNA*later* (Ambion) was added to preserve the RNA. Two biological replicates of WT and Δ*rclR* treated with HOCl or HOSCN, or untreated were collected for RNA purification. Enzymatic lysis and proteinase K digestion of bacteria followed by RNA purification was carried out using the RNeasy Protect Bacteria Mini Kit (QIAGEN). DNase treatment of RNA samples was performed using the TURBO DNA-free kit (Ambion). Removal of rRNA, preparation of cDNA libraries and sequencing using the Illumina NextSeq 500 system and a 75 bp read length was performed by vertis Biotechnologie AG. Reads were aligned to the *P. aeruginosa* UCBPP-PA14, complete genome (NCBI accession number: NC_008463.1) using bowtie2 and expression values in Reads Per Kilobase per Million mapped reads (RPKM) were calculated using FeatureCounts. Normalisation of the RPKM values against the OD_600_ of the samples prior to RNA extraction was performed and the values of the two biological replicates were averaged. The log2 fold change of the RPKM values of WT treated compared to WT untreated or WT treated compared to Δ*rclR* treated was calculated. Genes with a log2 fold change of >1.5 and <−1.5 were considered to be differentially expressed. The transcriptomic data have been deposited in the Gene Expression Omnibus database and are accessible through the GEO accession number: GSE124385.

### Quantitative real-time PCR

Cultures of PA14 WT were grown to mid-exponential phase in LB and treated with HOCl or HOSCN for 20 minutes prior to RNA extraction, this was performed as described for RNA_seq_. The cDNA library was prepared using the High-Capacity cDNA Reverse Transcription Kit (Applied Biosystems). Quantitative PCR was set up with Fast SYBR Green mix (Invitrogen) and using the Applied Biosystems™ 7500 Fast Real-Time PCR Instrument. Expression ratios were calculated comparing the expression of each gene in PA14 WT treated cultures to PA14 WT untreated, by the ΔΔCT method (95) and normalised to expression of *rpoD* (RNA polymerase sigma factor), which shows unchanged expression levels under the conditions tested. Primers used for qRT-PCR are listed in Table S1.

### β-Galactosidase assays

PA14 WT and Δ*rclR* strains containing *rclR-lacZ* and *rclX-lacZ* plasmids, and clinical CF isolates 1, 2 and 3 containing the *rclX-lacZ* plasmid were grown in LB medium with 100 μg/ml tetracycline overnight. Cultures were subcultured 1:50 (OD 0.05 ± 0.02) in LB medium in 2.2 ml 96-well deep well plates (VWR) and grown to exponential phase (3 hours) and were then either untreated or treated with HOCl (2.2 mM) or HOSCN (0.8 mM). Cultures were collected 30 minutes after treatment with HOCl or HOSCN. β-Galactosidase activity was assayed using the modified version (96) of the Miller method (97).

### Homology modelling and sequence analysis

The predicted structure of RclX was modelled with SWISS-MODEL (98) and illustrations of protein structures were prepared with EzMol (99). Amino acid sequences were aligned using Clustal Omega (100) and visualised with ESPript 3.0 (101).

## Acknowledgements

We thank Emily Calamita for her technical support in the construction of the Δ*pyeRM-xenB* mutant. The RNA_seq_ data have been deposited in the GEO database, accession number: GSE124385.

## FOOTNOTES

KVF was funded by a Medical Research Council Doctoral Training Grant and this work was also supported by the Cystic Fibrosis Trust Strategic Research Centre via a grant titled, Personalised approach to *Pseudomonas aeruginosa*.

The abbreviations used are:

CF: cystic fibrosis
HOCl: hypochlorous acid
HOSCN: hypothiocyanous acid
CFTR: <u>c</u>ystic <u>f</u>ibrosis <u>t</u>ransmembrane conductance <u>r</u>egulator
T3SS: type III secretion system
O_2_^−^: superoxide
H_2_O_2_: hydrogen peroxide
MPO: myeloperoxidase
Cl^−^: chloride
SCN^−^: thiocyanate
DUOX: dual oxidase
LPO: lactoperoxidase
HOBr: hypobromous acid
MFS: major facilitator superfamily
AhpD: alkylhydroperoxidase
TBH: *tert*-butyl hydroperoxide
NEM: *N*-ethylmaleimide
DEANO: diethylamine NONOate
LB: Luria-Bertani
ASM: artificial sputum media
GO: glucose oxidase
KSCN: potassium thiocyanate
TNB: 5-thio-2-nitrobenzoic acid
OD: optical density

